# Glycogen homeostasis and mtDNA expression require motor neuron to muscle TGFβ/Activin Signaling in *Drosophila*

**DOI:** 10.1101/2024.06.25.600699

**Authors:** Heidi Bretscher, Michael B. O’Connor

## Abstract

Maintaining metabolic homeostasis requires coordinated nutrient utilization between intracellular organelles and across multiple organ systems. Many organs rely heavily on mitochondria to generate (ATP) from glucose, or stored glycogen. Proteins required for ATP generation are encoded in both nuclear and mitochondrial DNA (mtDNA). We show that motoneuron to muscle signaling by the TGFβ/Activin family member Actβ positively regulates glycogen levels during *Drosophila* development. Remarkably, we find that levels of stored glycogen are unaffected by altering cytoplasmic glucose catabolism. Instead, Actβ loss reduces levels of mtDNA and nuclearly encoded genes required for mtDNA replication, transcription and translation. Direct RNAi mediated knockdown of these same nuclearly encoded mtDNA expression factors also results in decreased glycogen stores. Lastly, we find that expressing an activated form of the type I receptor Baboon in muscle restores both glycogen and mtDNA levels in *actβ* mutants, thereby confirming a direct link between Actβ signaling, glycogen homeostasis and mtDNA expression.

**Key Points:**

- The Drosophila TGFβ family member Actβ signals from motor neuron to muscle positively regulating glycogen levels
- Actβ positively regulates nuclearly encoded factors required for mtDNA expression
- Genes involved in mtDNA expression directly regulate glycogen stores
- Expressing an activated receptor in muscle restores glycogen and mtDNA in *actβ* mutants

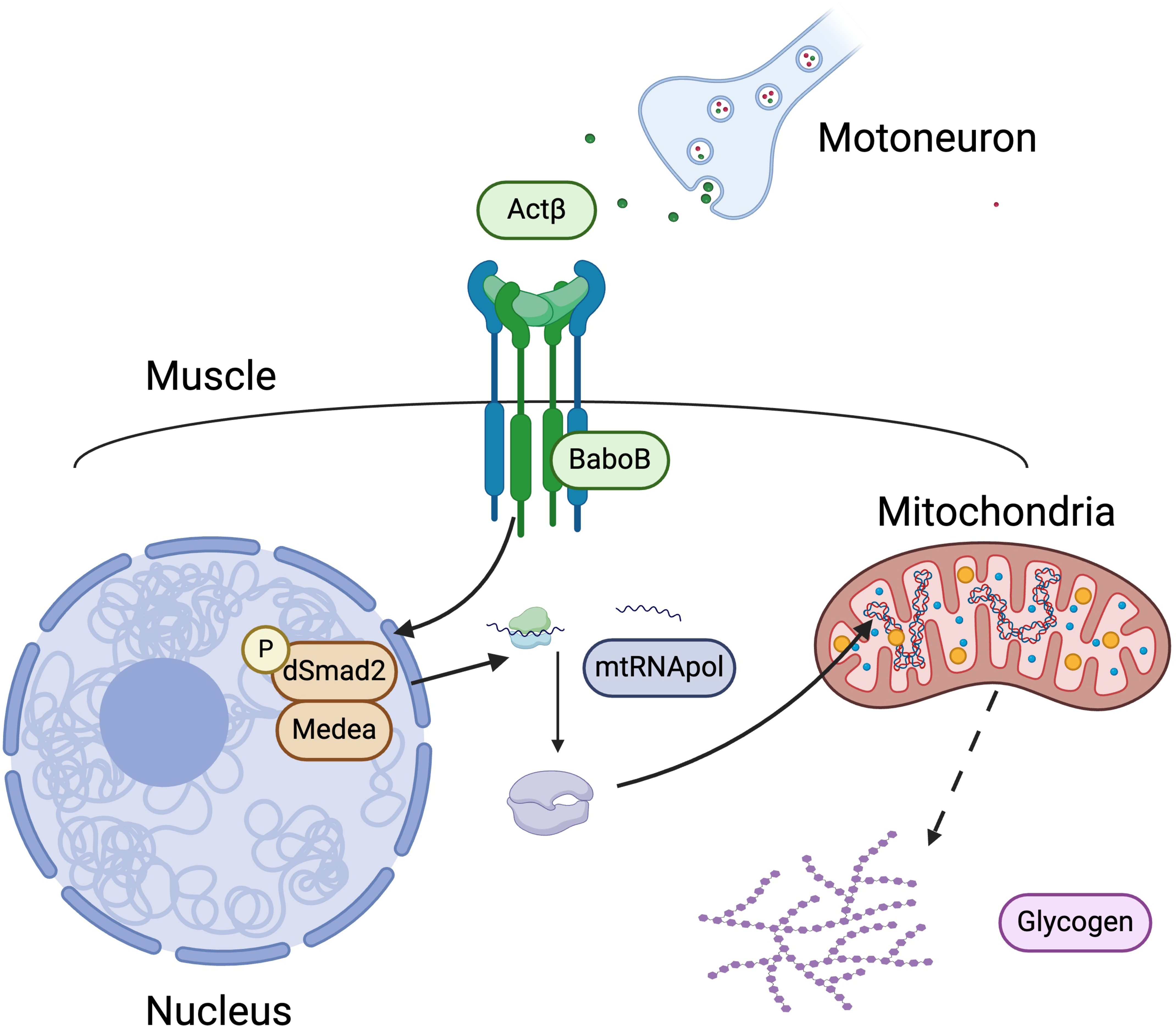

## Introduction

To maintain metabolic homeostasis, an organism must coordinate energy absorption, storage, and utilization between numerous organelles within a cell and across multiple organ systems. Central to energy production and metabolic homeostasis are mitochondria, which serve as the main source of ATP in most cell types. The energy required for ATP synthesis is generated during oxidative phosphorylation which consists of high energy electron transfer through the electron transport chain (ETC). Unlike most organelles, mitochondria have their own DNA. The mitochondrial genome is a maternally inherited circular plasmid and each individual mitochondria has multiple copies of its genome^1–3^. Mitochondrial DNA (mtDNA) is well conserved across metazoans and contains the sequences for 37 genes including 13 subunits of the ETC^2–5^, the exception being nematodes in which one subunit has been lost^6^. These 13 subunits make up parts of complexes I, III, IV and V, with the remaining subunits being encoded in the nuclear DNA. The mitochondrial genome does not code for transcriptional machinery, and thus is dependent on nuclear encoded machinery for replication, transcription and translation and thereby expression of its genes^1^. The expression of mitochondrially encoded ETC subunits and nuclearly encoded ETC subunits must be coordinated so that the ETC complexes come together in the correct stoichiometric ratios. This is carried out by a host of different nuclearly encoded products, most notable among them the mtRNA polymerase (PolrMT), the mtDNA polymerase (PolG1) and the mtDNA helicase (Twinkle)^2,4,5,7^. Failure to properly regulate the expression of mitochondrially encoded genes results in changes in energy metabolism and many distinct diseases^8–10^.

Mitochondrial diseases produce a diverse class of symptoms that primarily affect organs with high metabolic needs including muscle and the nervous system^1,3,10,11^. Approximately 400 genes, both mitochondrial and nuclear in origin, have been associated with mitochondrial diseases, and about 25% of these genes affect factors involved in mtDNA replication and transcription^3^. More than 300 disease causing mutations have been reported within PolG1 and Twinkle alone^3,11^. While symptoms vary across patients, some of the most common and unique symptoms are short stature, exercise intolerance, mitochondrial encephalomyopathy with lactic acidosis and stroke-like episodes (MELAS), and progressive external ophthalmoplegia (PEO) characterized by limited eye movements and ocular muscle weakness^3,11^. Interestingly, only eight individuals have been identified with PolrMT mutations, and these patients exhibit decreased expression of mitochondrially encoded genes, accumulation of abnormally sized mitochondria and ragged muscle fibers^10^. These diseases demonstrate the importance of regulating mtDNA expression in human health.

One of the important environmentally derived nutrients used as a substrate for mitochondrial ATP production in many organisms is glucose. Glucose can be stored in long branched chains known as glycogen which serves as an energy source during periods of high metabolic demand or when environmental nutrients are scarce^12^. In humans the main sites of glycogen stores are liver and muscle^12^ which are crucial for health and development. Glycogen storage diseases (GSDs) are a large class of diseases consisting of at least eight distinct types that affect glycogen homeostasis. GSDs range in severity and affect different tissues resulting in diverse phenotypes including exercise intolerance, altered metabolism, stunted growth and development and neurological abnormalities^12,13^. One of the more severe forms, GSD Type 0, results in a complete lack of glycogen and even with medical intervention to help compensate for this deficiency, cardiomyopathy and even death can occur demonstrating the importance of glycogen stores in development and survival^12,13^.

Elevated levels of glycogen have negative consequences as well. Patients with GSD type 1 are unable to mobilize glycogen which, when left untreated, can result in death during childhood from tissue damage caused by excess glycogen^13^. In addition to GSDs, excess glycogen accumulation in liver can drive tumorigenesis^14^ and high levels of glycogen in neurons is responsible for the neurodegeneration observed in Lafora disease^15^. Despite the importance of regulating glycogen levels, little is known about how basic metabolic pathways contribute to glycogen homeostasis during development.

*Drosophila* provides a unique model to study metabolic homeostasis during development. The life cycle of *Drosophila* consists of a 4-5 day larval stage in which animals increase in mass by approximately 200-fold. This is followed by a non-feeding pupal stage lasting 5 days during which extensive energy dependent remodeling occurs. Inadequate energy stores prior to pupation result in either failure to pupate or lethality during pupation^16^. During development, glycogen is stored in both the muscle and fat body (analogous to liver and adipose tissue)^17^. Animals unable to synthesize glycogen (*glyS* mutants) or animals unable to mobilize stored glycogen (*glyP* mutants) die at high rates during the larval stage,^18^ underscoring the importance of glycogen homeostasis for development and survival. Rare adult survivors lacking glycogen show decreased climbing ability and flight performance^18^. In addition to being required for survival, glycogen also serves as an important energy source during times of stress. In response to starvation, glycogen is rapidly mobilized from the fat body of young larvae^19^, and muscle^20^. Additionally, muscle glycogen plays an important role in surviving parasitic wasp infection^21^.

Coordination of energy absorption, storage and utilization is carried out in part by cell-signaling pathways. While cell-signaling pathways stimulated by insulin^22,23 24–26^ and glucagon^27–30^ have well established central roles in regulating metabolic homeostasis, it has recently been appreciated that other cell signaling pathways traditionally known for their role in development and patterning, such as Wnt^31^, hedgehog^32^, JAK/STAT^21,33,34^ and TGFβ^35–37^ are also crucial to metabolic homeostasis.

The TGFβ superfamily consists of two branches whose members form the BMP and TGFβ/Activin subfamilies of ligands. In *Drosophila* the TGFβ/Activin branch includes three ligands, Myoglianin (Myo), Activin-β (Actβ) and Dawdle (Daw)^38^. All three ligands signal through the same type I receptor, Baboon, in conjunction with a type II receptor, Punt or Wit. Baboon has three distinct splice isoforms, termed Babo A, Babo B and Babo C, that differ only in exon 4, which encodes the ligand binding domain. This difference enables each ligand to potentially signal through a single splice isoform^39^. Formation of the ligand receptor complex stimulates phosphorylation of the receptor-Smad, dSmad2/Smox, which translocates to the nucleus as a complex with the co-Smad Medea where it serves as a transcriptional transducer^38^. Each ligand and splice isoform have a unique expression pattern enabling an individual response of each tissue to a different complement of ligands depending on the origin of the ligand and the receptor profile of the receiving tissue. Such a mechanism enables this branch of the superfamily to control different sets of developmental and metabolic genes and processes in a tissue specific manner^35,40–45^.

In this report, we demonstrate that the TGFβ/Activin ligand, Actβ positively regulates both glycogen levels and nuclearly encoded transcriptional machinery required to maintain the mtDNA genome. Loss of Actβ results in decreased glycogen, decreased mtDNA content and decreased mitochondrially encoded gene transcription. These phenotypes can be rescued by expressing an active form of the type I receptor, Baboon, in muscle. A candidate screen demonstrates that maintenance of glycogen levels in wild-type animals is robust and independent of disruptions in cytoplasmic glucose catabolism. Instead, glycogen homeostasis during organismal development is regulated by mitochondria and factors involved in mtDNA expression demonstrating a direct link between nuclearly encoded mitochondrial transcriptional machinery and carbohydrate metabolism.

## Results

### Actβ positively regulates glycogen, but not lipid, levels in both body wall muscle and fat body

We have previously shown that loss of the TGFβ/Activin ligand Actβ results in small pharate lethal pupae despite normal feeding behavior ^45^, which led us to ask whether Actβ is required to maintain metabolic homeostasis during development. We chose to look at early L3, pre-critical weight larvae, 68-72 hours after egg lay (AEL), as at this developmental time point animals grow rapidly and respond to their nutritional environment by storing and mobilizing nutrients according to metabolic needs^46^. We began by measuring glycogen levels and found that two independent null alleles of Actβ, *act^80^ and act^4E^*, both have severely depleted glycogen stores (Fig 1A). Homozygous *actβ* mutants have 70% less stored glycogen when compared to control larvae (*w^1118^*), and heterozygous *actβ* larvae (act*^80^*/+) have 20% less than controls (Fig 1A), indicating that Actβ is a strong positive regulator of glycogen stores. In *Drosophila* glycogen is stored in both the body wall/skeletal muscle and fat body (analogous to the liver and adipose tissue), so we next examined whether Actβ regulates glycogen levels in these tissues. Since glycogen is the most abundant carbohydrate in *Drosophila* muscle and fat body^19^, we used Periodic Acid Schiff’s Reagent to stain for glycogen and found that *actβ* mutants have less glycogen in both muscle (Fig 1B,C) and fat body (Fig 1D,E). One possible reason for decreased glycogen levels is increased glucose levels. At this point in development the level of organismal free glucose is very low, however we were able to detect a small amount of glucose and found that Actβ is also a positive regulator of glucose levels (Fig S1A). To explore the possibility that glucose uptake may be decreased in *actβ* mutants, we used a fluorescent glucose analog, 2-NBDG, to monitor glucose uptake in the muscle *ex-vivo*. We found that glucose uptake varies considerably between individual muscle segments but found no difference between genotypes (Fig S1B-D).

**Figure 1:**
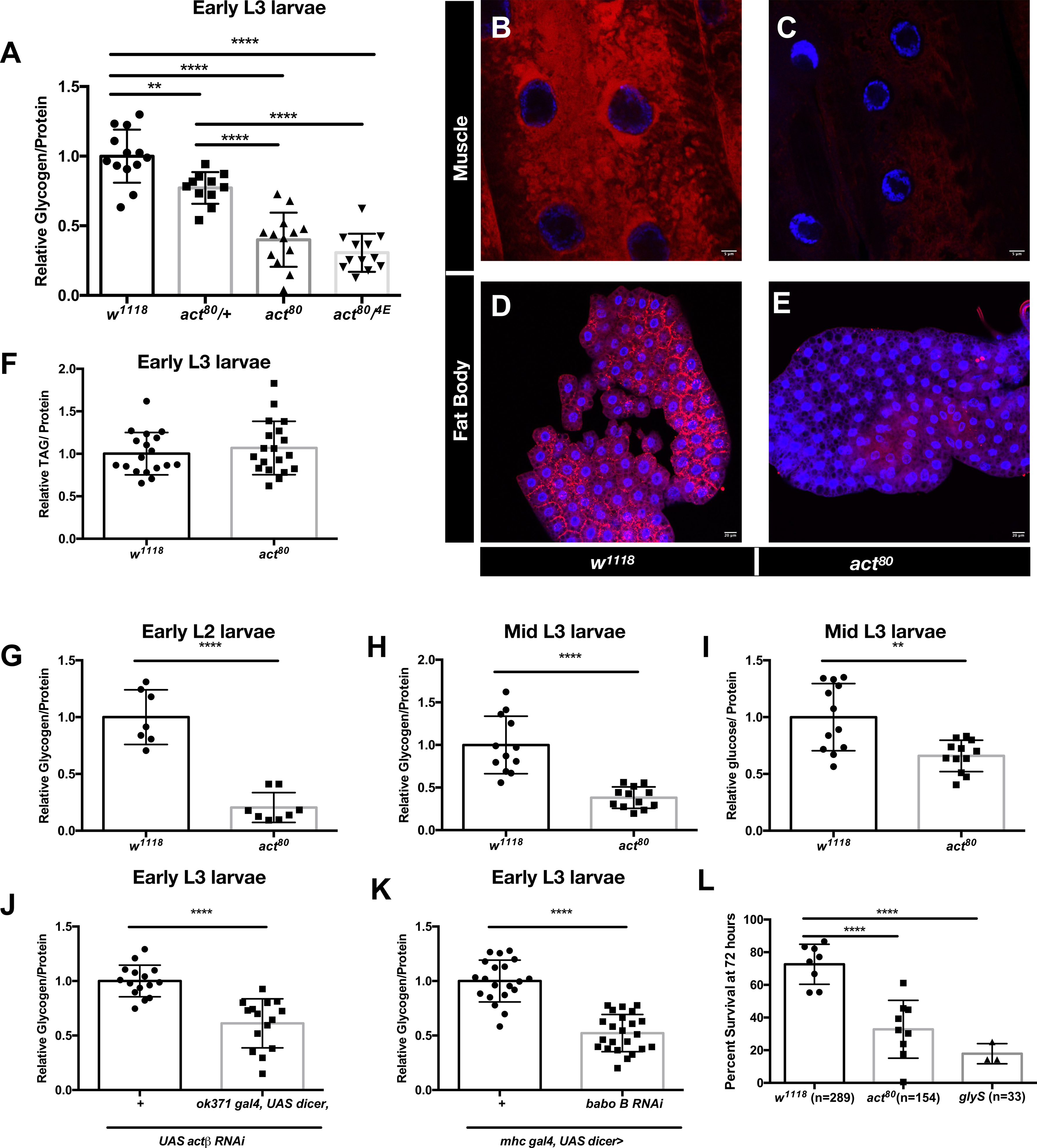
Actβ positively regulates glycogen levels during *Drosophila* larval development. A) Quantification of total glycogen/protein in early L3 larvae of control (*w^1118^*), heterozygotes *actβ* mutants (*act^80^/+*), homozygous null *actβ* mutants (*act^80^*) and transheterozygous null *actβ* mutants (*act^80/4E^)*. Periodic Acid/SchiB’s Reagent staining of skeletal muscle of B) control (*w^1118^*) C) *actβ* mutants (*act^80^*) and fat body of D) control (*w^1118^*) E) *actβ* mutants (*act^80^*). All animals are early L3 larvae. F) Quantification of total lipids per protein in early L3 larvae of control (*w^1118^*) and homozygous null *actβ* mutants (*act^80^*). G) Quantification of total glycogen per protein in early L2 larvae (∼54 hours AEL) of control (*w^1118^*) and homozygous null *actβ* mutants (*act^80^*). H) Quantification of total glycogen per protein in mid-L3 larvae (∼96 hours AEL) of control (*w^1118^*) and homozygous null *actβ* mutants (*act^80^*). I) Quantification of total free glucose per protein in mid-L3 larvae (∼96 hours AEL) of control (*w^1118^*) and homozygous null *actβ* mutants (*act^80^*). J) Quantification of total glycogen/protein in early L3 larvae of control (*+>UAS actβ RNAi)* and motor neuron specific knockdown of *actβ (ok371 gal4 > UAS dicer, UAS actβ RNAi*). K) Quantification of total glycogen/protein in early L3 larvae of control (*mhc gal4 > +)* and muscle specific knock down of *babo B* ((*mhc gal4 > UAS babo B)*. In all panels each data point represents 5 individual animals except panel G in which each data point represents 15 animals. All experiments were repeated a minimum of 3 times. L) Percent of early L3 larvae of control (*w^1118^*), homozygous null *actβ* mutants (*act^80^*) and homozygous null *glycogen synthase* mutants *(glyS)* surviving a 72 hour starvation period. Each data point represents an individual experiment and total animals tested is listed under genotype. All animals were raised until designated time point on apple juice plates supplemented with yeast paste (60% w/v in water). * = p<0.05, **=p<0.01, ***=p<0.001, ****=p<0.0001

In addition to storing nutrients as glycogen, *Drosophila* also stores lipids as nutrients. No difference in amount of stored lipids in *actβ* mutants compared to controls was found (Fig 1F) demonstrating that Actβ positively regulates muscle and fat body glycogen levels independent of glucose uptake but is dispensable for lipid homeostasis.

We next examined the role of Actβ in regulating glycogen levels throughout the *Drosophila* larval life cycle. Early in larval development (the L1/L2 transition, about 54 hours AEL, Fig 1G), and later in larval development (during mid-L3 when animals are post-critical weight, ∼96 hours AEL, Fig 1H), *actβ* mutants also have significantly less glycogen than control animals. In addition, by mid-L3 stage we detected a significant amount of free glucose in whole organisms and found that Actβ is also a positive regulator of free glucose (Fig 1I).

Since Actβ is a secreted ligand, we next sought to determine the tissue(s) in which signal reception is required for maintaining glycogen homeostasis. We have previously shown that Actβ is expressed in motor neurons which synapse on the larval body wall/skeletal muscle^40,45^. Knockdown of *actβ* in motor neurons of early L3 larvae results in decreased organismal glycogen levels (Fig 1J). To determine whether the muscle is an important tissue in receiving the signal we knocked down Babo B, the Actβ specific type 1 receptor isoform^47^, in muscle and again found a significant decrease in organismal glycogen levels (Fig 1K). Thus, we conclude that motor neuron derived Actβ signals through Baboon B in the body wall/ skeletal muscle to directly regulate the amount of stored glycogen. In the remaining sections, we chose to focus primarily on the muscle as this is the main source of stored glycogen at the early L3 larval stage^19^.

### Actβ and glycogen stores are required for optimal starvation survival

Given that *actβ* mutants have decreased glycogen levels, we assessed whether they are starvation sensitive at the early pre-critical weight L3 stage since at this developmental time point larvae mobilize stored metabolites in response to starvation and die from starvation rather than accelerate pupation^16^. We found that while ∼75% of control larvae survive a 72-hour starvation period, only 32% of *actβ* mutants survive 72 hours of nutrient deprivation (Fig 1L). To determine whether glycogen is required for starvation survival we next investigated whether a null mutation in glycogen synthase (GlyS)^18,19^, which is required to synthesize glycogen^18,19^, also renders animals starvation sensitive. Only 18% of *glyS* mutant larvae survive a 72-hour starvation (Fig 1L) demonstrating the importance of glycogen stores in the ability to withstand nutrient deprivation. Therefore, we hypothesize that decreased glycogen contributes to the starvation sensitivity of *actβ* mutants.

### Blocking glycogen catabolism does not restore glycogen level in *actβ* mutants

Glycogen is made up of long branched chains of glucose, which is synthesized by Glycogen Synthase (GlyS) and broken down by Glycogen Phosphorylase (GlyP) (Fig 2A). One possible explanation for decreased glycogen levels in *actβ* mutants is increased glycogen catabolism. To investigate this possibility, we introduced the null *glyP*^18,19^ mutation into the *actβ* mutant background and investigated whether this restores glycogen levels. Although we saw a significant increase in glycogen levels (Fig 2B), the *glyP;;actβ* double mutant still has significantly less glycogen than the *glyP* mutant alone (Fig 2B), demonstrating that increased GlyP activity does not account for the observed reduced glycogen levels in *actβ* mutants. Moreover, the fact that the *glyP;;actβ* double mutant has more glycogen than *actβ* single mutants (Fig 2B) indicates that Actβ is not required for glycogen synthesis. In support of this notion, overexpression of GlyS in the muscle of *actβ* mutants also fails to restore glycogen levels (Fig S1A). Finally, we did not observe any differences in mRNA levels of either *glyS* or *glyP* (Fig S1B) in *actβ* mutants.

**Figure 2:**
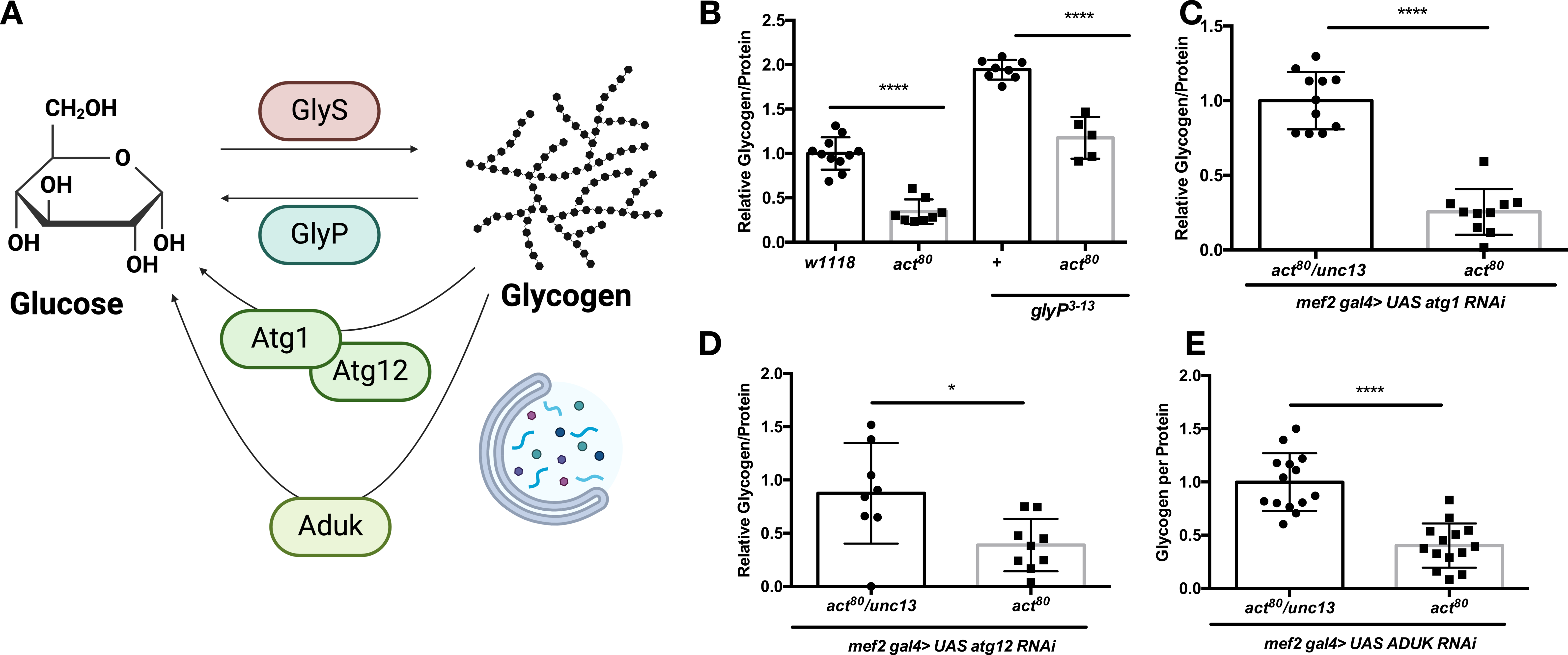
Blocking glycogen catabolism fails to restore glycogen levels in *actβ* mutants. A) Schematic representing glycogen synthesis and glycogen catabolism. GlyS=glycogen synthase, GlyP= glycogen phosphorylase, Atg1=autophagy related 1, Atg12=autophagy related 12. Glycogen is synthesized by GlyS, and can be broken down via GlyP or via canonical (Atg1/12 dependent) or non-cannonical (Aduk dependent) autophagy. Quantification of glycogen/protein in B) control (*w^1118^*), homozygous null *actβ* mutants (*act^80^*), homozygous null *glyP* mutants (*glyP^3-13^*) and homozygous null *actβ, glyP* double mutants (*glyP^3-13^;;act^80^*) C) muscle specific knockdown of *atg1* (*mef2gal4>UAS atg1* RNAi) in *actβ* heterozygotes (*act^80^/unc13*) and *actβ* homozygous mutants (*act^80^*) D) muscle specific knockdown of *atg12* (*mef2gal4>UAS atg12* RNAi) in *actβ* heterozygotes (*act^80^/unc13*) and *actβ* homozygous mutants (*act^80^*)) muscle specific knockdown of *aduk* (*mef2gal4>UAS aduk* RNAi) in *actβ* heterozygotes (*act^80^/unc13*) and *actβ* homozygous mutants (*act^80^*). All animals are early L3 raised on apple juice plates supplemented with yeast paste (60% w/v in water). In all panels each data point represents 5 individual animals. All experiments were repeated a minimum of 3 times. * = p<0.05, **=p<0.01, ***=p<0.001, ****=p<0.0001

While autophagy is not thought to be a common source of glycogen breakdown in mammals^12^, it has been shown to be a significant source of glycogen catabolism upon starvation in flies ^20^. Therefore, we next attempted to restore glycogen levels in *actβ* mutants by inhibiting autophagy in the muscle. To control for potential background effects in RNAi experiments we chose to compare *actβ* heterozygotes (*act^80^/unc13*) to *actβ* homozygous mutants (*act^80^*), so that all offspring had the same parents. While we have shown that Actβ is partially haploinsufficient when it comes to regulating glycogen levels (Fig 1A), we reasoned that the disparity in glycogen levels between heterozygotes and homozygotes should still be sufficient to determine if there is a differential response to knockdown of autophagy. We expressed RNAi in muscle targeting *atg1*^48^, an important regulator of autophagy and important in formation of the pre-autophagosome complex, and *atg12*^48^, a downstream protein also involved in autophagosome formation, but neither knockdown restored glycogen levels in *actβ* mutants (Fig 2C,D). Similarly, inhibiting non-canonical, Atg1 independent autophagy^49^, by knocking down *aduk* in the muscle also failed to eliminate glycogen deficits in *actβ* mutants (Fig 2E). Therefore, we conclude that increased glycogen catabolism by either GlyP or autophagy does not explain reduced glycogen levels in *actβ* mutants.

Since insulin signaling is required for glycogen synthesis in *Drosophila*^19^, we investigated whether decreased insulin signaling could account for decreased glycogen levels upon loss of Actβ. To increase insulin signaling we overexpressed two proteins downstream of insulin activation, Pdk1^50^ (Fig S2C) and a constitutively active form of Akt, AktT342D^51^ (Fig S2D) in the muscle, however neither of these manipulations restores glycogen levels in *actβ* mutants. We also increased insulin signaling by overexpressing Dilp5^52^, a *Drosophila* Insulin-like peptide, in the insulin producing cells and again this did not eliminate the glycogen deficiency in *actβ* mutants (Fig S2E). A second pathway with numerous established roles in metabolic homeostasis and nutrient mobilization and breakdown is glucagon signaling^28–30,35^. To explore the possibility that the effect we were seeing on glycogen homeostasis was due to mis-regulated glucagon signaling, we introduced a null mutation in the receptor for Akh (the *Drosophila* glucagon homolog), AkhR^53^ into the *actβ* mutant background. This did not alter the glycogen depletion in *actβ* mutants (Fig S2F). Finally, we investigated the possibility that the diet we were using did not contain enough glucose to support glycogen synthesis in *actβ* mutants. Therefore, animals were raised on a high sugar diet, and yet again, the glycogen deficiency in *actβ* mutants persisted (Fig S1G). Taken together, we conclude that Actβ’s role in regulating glycogen homeostasis is independent of basic glycogen synthesis and catabolism as well as processes with well-known roles in metabolic homeostasis.

### A screen identifies mitochondria as pivotal regulators of glycogen metabolism

To gain an insight into metabolic processes that regulate glycogen homeostasis during development, we undertook a candidate screen to identify basic steps in glucose catabolism and energy production that affect levels of organismal glycogen in wild-type animals. We again chose to look at pre-CW L3 larvae and focused on the muscle, since this tissue is an important source of stored glycogen^19^. The results of this screen are summarized in Fig 3A, in which loss of enzymes in blue have no effect on glycogen homeostasis, loss of enzymes in pink have a modest effect on glycogen levels and loss of enzymes in red are strong regulators of glycogen homeostasis. One of the main pathways in which glucose is broken down to yield cellular energy is glycolysis. Since glycogen is synthesized from glucose, we investigated whether blocking glycolysis alters glycogen homeostasis. RNAi in the muscle directed at two different glycolytic genes encoding *phospho-6-glucose isomerase (pgi)*^54^ and *phosphofructokinase (pfk)*^54^ does not produce a change in glycogen levels (Fig 3B). To explore the possibility that glycolysis was being upregulated in other tissues to compensate for loss in the muscle, we also measured levels of glycogen in two glycolytic pathway mutants, *enolase (eno)*^55^ (Fig 3C) and *pgi*^55^ (Fig 3D), but again this did not affect glycogen levels. The process of gluconeogenesis, which maintains glucose levels during starvation, relies on reversing glycolysis and is dependent on Pepck to regenerate the glycolytic intermediate phosphoenolpyruvate. Neither knockdown of *pepck*^56^ in muscle (Fig 3B), nor a *pepck* mutant (Fig 3E) show any alteration in glycogen levels.

**Figure 3:**
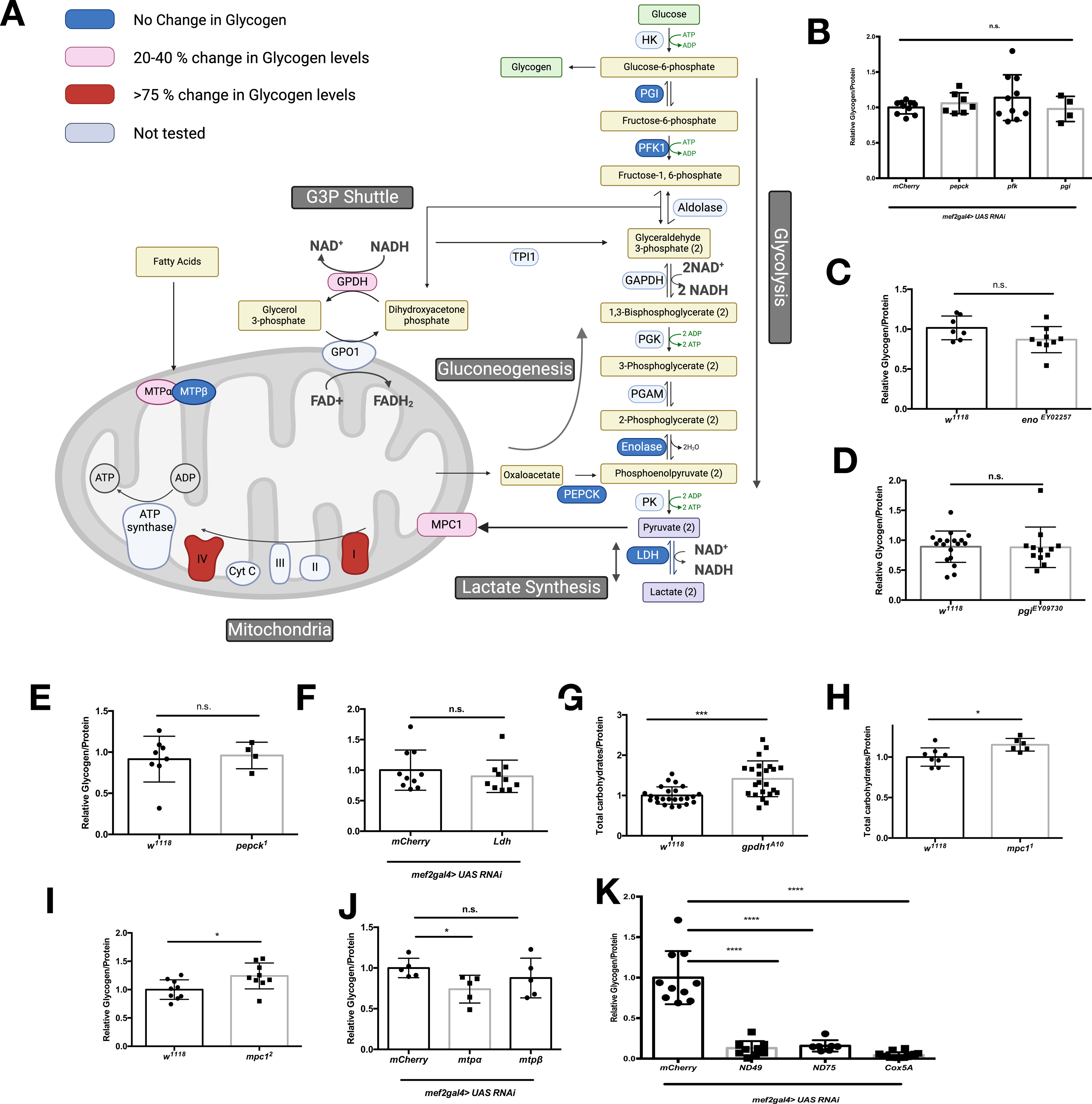
A screen of basic metabolic process identifies mitochondria as central regulators of glycogen homeostasis during *Drosophila* larval development. a) Schematic representing metabolic process tested in screen. Muscle specific knockdown (*mef2 gal4 >RNAi*) and/or mutants of enzymes depicted in blue resulted in no change in glycogen homeostasis, enzymes in pink resulted in a 20-40% change in glycogen homeostasis and enzymes in red produced a greater than 75% change in glycogen level. Enzymes in gray were not tested. b-f and i-k) quantification of glycogen per protein in designated genotype. g,h) quantification of total carbohydrates (free glucose + glycogen) per protein in designated genotype. All animals are early L3 raised on apple juice plates supplemented with yeast paste (60% w/v in water). In all panels each data point represents 5 individual animals. All experiments were repeated a minimum of 2 times. * = p<0.05, **=p<0.01, ***=p<0.001, ****=p<0.0001

We were initially surprised that blocking glycolysis failed to alter glycogen levels, however, we reasoned that this may be because some glycolytic enzymes are semi-redundant with enzymes in other pathways. Therefore, we next targeted pathways important in maintaining redox balance and regenerating cytoplasmic NAD+ allowing constant flux through glycolysis. The main pathway used by the muscle to maintain cytoplasmic redox homeostasis is the lactic acid pathway catalyzed by Lactate Dehydrogenase (Ldh)^57^ (Fig 3A); however, knockdown of *Ldh* in muscle has no effect on organismal glycogen (Fig 3F). A second pathway that regenerates NAD+ for glycolysis, is the glycerol-3-phosphate shunt (Fig 3A). This pathway, catalyzed by Glycerol-3-phosphate dehydrogenase 1 (Gpdh1), regenerates cytoplasmic NAD+ and is coupled to a mitochondrial reaction, catalyzed by Gpo1, which generates mitochondrial FADH_2_. When looking at levels of glycogen in *gpdh1* mutants^57^ we found a combination of increased glucose and increased glycogen. Therefore, we decided to look instead at total carbohydrates and found that *gpdh1* mutants have a 40% increase in total carbohydrates (Fig 3G) in agreement with previously published data^57^. Since the glycerol-3-phosphate shunt involves the mitochondria and glucose derivatives being used to generate energy in mitochondria, this gave us a hint that mitochondria, but not cytoplasmic glucose catabolism, may regulate glycogen homeostasis.

A second glucose derivative used for mitochondrial energy generation is pyruvate, which is generated by glycolysis and enters the mitochondria via the Mitochondrial Pyruvate Carrier (*mpc1).* Two independent hypomorphic alleles of *mpc1*^58^, have ∼20-25% increased total carbohydrate levels (again we observed some increase in glucose and some increase in glycogen with the *mpc1^1^* allele) (Fig 3H,I), in agreement with previously published data^58^. Having seen that inhibiting the entry of glucose derivatives into mitochondria results in altered glycogen homeostasis, we next examined whether inhibiting fatty acids from being used as a mitochondrial substrate would alter glycogen homeostasis. Blocking mitochondrial fatty acid β-oxidation, by knocking down *mtpα* in muscle produces a 25% reduction in glycogen levels (Fig 3J). Taken together, our results demonstrate that limiting mitochondrial substrates, but not manipulating individual steps in cytoplasmic glucose catabolism alter glycogen homeostasis during *Drosophila* larval development.

To further explore the role of mitochondrial energy generation in regulating glycogen levels, we next targeted oxidative phosphorylation via knock down of components of Complex 1 (*ND-75 and ND49*) and Complex IV (*Cox5A)*. All three of these knockdowns result in greater than 75% reduction in glycogen levels (Fig 3K), with no corresponding increase in glucose levels. **I made a graph for this but not sure where to put it/ if we need it.** Thus, we conclude that mitochondria are crucial for glycogen homeostasis during *Drosophila* larval development.

### Blocking entry of glucose derivatives into mitochondria does not restore glycogen levels in *actβ* mutants

Having seen that inhibiting glucose derivatives from being used for mitochondrial energy generation in otherwise wildtype animals mildly increases glycogen levels, we wondered whether Actβ may regulate glycogen levels via altering the flux of glucose derivatives into mitochondria. Even though we saw no change in glycogen levels in glycolytic or gluconeogenic mutants, we explored the possibility that increased glycolysis/gluconeogenesis was responsible for decreased glycogen in *actβ* mutants. We first introduced mutants in two glycolytic genes, *pgi* (Fig S3A) and *eno* (Fig S3B) into the *actβ* mutant background and found that this did not restore glycogen levels. Similarly, inhibiting gluconeogenesis with a mutation in Pepck (Fig S3C) does not rescue the glycogen deficit observed in *actβ* mutants.

Since our screen demonstrated that pyruvate entry into mitochondria regulates glycogen homeostasis, we assessed whether inhibiting this step could restore glycogen levels in *actβ* mutants. As above, due to concerns over different background, we chose to compare *actβ* homozygous mutants (*act*^80^) to heterozygotes from the same stock (*act*^80^*/unc13*) so that all animals had the same parents. Two independent hypomorphic mutations in *mpc1*^58^ fail to restore glycogen in *actβ* mutants (Fig S 3D,E). Similarly inhibiting either shuttle required to maintain cytoplasmic redox balance, the glycerol-3-phosphate shuttle (Fig S3F) or lactate production (Fig S3G), did not rescue glycogen levels in *actβ* mutants. Taken together, these results demonstrate that Actβ does not regulate glycogen homeostasis via limiting mitochondrial substrate availability or increasing glycolytic flux.

### Actβ positively regulates mitochondrial DNA content, but not overall mitochondrial protein abundance

Having identified mitochondria as important regulators of glycogen homeostasis, we next investigated whether Actβ regulates mitochondrial homeostasis. Since mitochondria have their own DNA, the ratio of total genomic mitochondrial DNA (mtDNA) to total genomic nuclear DNA (nDNA) can be used as a readout for mitochondrial content. To control for any differences in maternally inherited mtDNA, we compared *actβ* mutants (*act^80^*) to *unc13/+* (generated by crossing *act^80^/unc13-gfp* females to *w^1118^* and selecting for GFP) or *actβ* heterozygotes (*act^80^/+* generated by crossing *act^80^/unc13-gfp* females to *w^1118^* and selecting against GFP) so that all comparisons have the same maternal mtDNA. When comparing *actβ* mutants to *unc13/+* we find a 60% reduction in mtDNA content (Fig 4A) and comparing *actβ* mutants to heterozygotes results in a 35% reduction of mtDNA (Fig 4B). Therefore, we conclude that Actβ positively regulates mtDNA. To test whether Actβ positively regulates mitochondrial protein levels, we analyzed the protein abundance of a nuclearly encoded component of the ATPSynthase (Complex V), (ATP5A) and were surprised to see no difference in ATP5A protein levels in *actβ* mutants (Fig 4C,D). To ensure this was not limited to a single subunit of the ETC, we also examined the abundance of ND30/NDFSU3, a nuclearly encoded subunit of complex I. Again, we did not observe any changes in protein abundance of nuclearly encoded ND-30 (Fig S4A,B). Since our main tissue of interest is the muscle, we stained muscles with an antibody directed at ATP5A and measured the area occupied by mitochondria in the subsarcolemmal plane and observed no difference in total area occupied by mitochondria (Fig 4E-G) in agreement with our western blot analysis (Fig 4C,D). While we did not see a difference in mitochondrial area, we did notice a difference in morphology. Loss of Actβ leads to a more condensed fragmented mitochondrial network with fewer distinct uninterrupted rod/tubular structures (Fig 4 H-I). To quantify this effect, we selected a 200 pixel by 200 pixel area in muscle 6 or 7 and measured the length of the five longest distinct rod structure (Fig S4C-E) and found that it is shorter in *actβ* mutants (Figure 4J) demonstrating that while Actβ does not regulate overall mitochondrial protein content, it is involved in regulating mtDNA levels and mitochondrial morphology.

**Figure 4:**
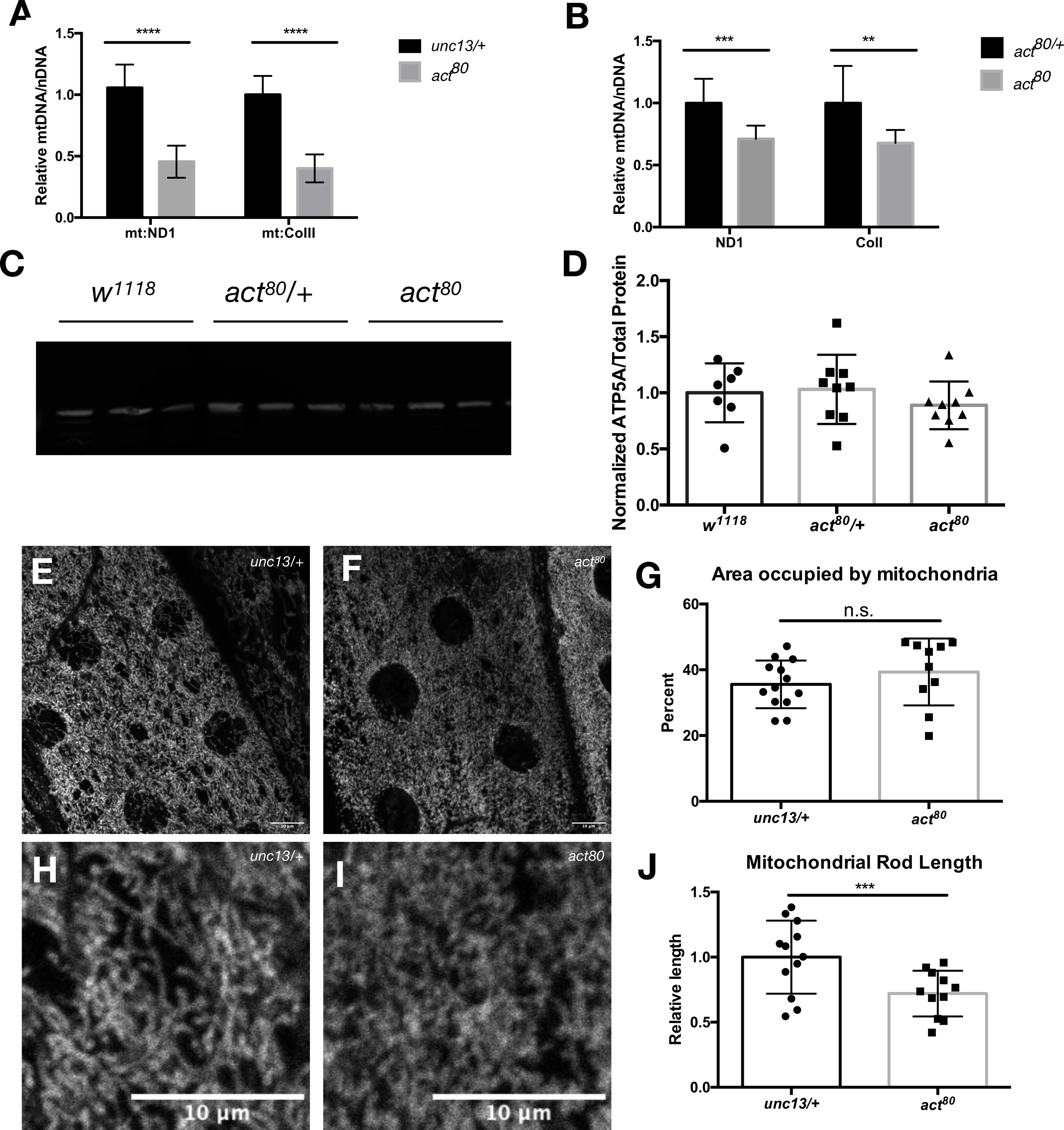
Actβ positively regulates mitochondrial DNA (mtDNA) content but not nuclearly encoded mitochondrial protein levels. q-PCR Quantification of the ratio of mtDNA (*mt:ND1* and *mt:CoI*) to nDNA (*rpl23*) in early L3 larvae of A) control (*unc13/+*) and *actβ* homozygous mutants (*act^80^*) or B) *actβ* heterozygotes (*act^80^/+)* and *actβ* homozygous mutants (*act^80^*). N=7 biological replicates consisting of 5 animals each for all conditions. C) Western blot showing expression level of nuclearly encoded mitochondrial protein ATP5A/blw in early L3 larvae of control (*w^1118^*), *actβ* heterozygotes (*act^80^/+*), and homozygous null *actβ* mutants (*act^80^*). All lanes represent 5 animals and 20μg of protein was loaded. D) Quantification of ATP5A expression levels across 3 independent western blots. Early L3 larvae skeletal muscle stained with ATP5A to mark mitochondria of E) control (*unc13/+*) and F) homozygous null) *actβ* mutants (*act^80^*. G) Total area occupied by mitochondria in skeletal muscle. Each data point represents one animal. Early L3 larvae skeletal muscle stained with ATP5A to show mitochondrial morphology of H) control (*unc13/+*) and iI homozygous null *actβ* mutants (*act^80^*). j) Quantification of mitochondrial rod length. * = p<0.05, **=p<0.01, ***=p<0.001, ****=p<0.0001

### Actβ positively regulates mtDNA expression factors and maintains the balance of nuclearly and mitochondrially encoded OXPHOS subunits

Having found a role for Actβ in regulating mtDNA, but not overall mitochondrial protein, we took a transcriptomics approach to determine possible causes for this phenotype. We find that Actβ positively regulates expression level of several nuclearly encoded factors involved in mtDNA replication, transcription and translation (collectively referred to as mtDNA expression) such as mtRNA polymerase (mtRNApol, PolrMT), which is the sole mitochondrial RNA polymerase in flies,^2,^ mtDNA helicase (Twinkle), a second mitochondrial helicase^2^, Suv3, (Fig 5A) and the mitochondrial translation elongation factor mEFG1. Other factors positively regulated by Actβ involved in mtDNA expression are shown in Fig 5A. This data provides an explanation for our findings in Fig 4 where we show Actβ is a regulator of mtDNA levels. Since mtDNA is reliant on these nuclearly encoded factors for expression, then it is likely that mitochondrially encoded, but not nuclearly encoded subunits of the ETC may be down regulated in *actβ* mutants. Transcriptome analysis reveals that expression of mitochondrially, but not nuclearly, encoded subunits of Complex I (the largest complex in the ETC) are down regulated in *actβ* mutants (Fig 5B). This explains why we did not observe any differences in overall mitochondrial protein levels despite differences in mtDNA levels (Fig 4A-G). Given these results we reasoned that we should be able to visualize a decrease in mtDNA and a decrease in the ratio of mtDNA to nuclearly encoded mitochondrial protein in muscle. We took advantage of a very sensitive DNA dye, picogreen, which can be used to visualize and quantify mtDNA^59–61^. We found that the signal intensity of picogreen in the muscle was significantly decreased in *actβ* mutants when compared to controls (Fig 5C,D first panel, Fig5E). In addition, co-staining with nuclearly encoded inner mitochondrial membrane protein ATP5A, we found that the ratio of mtDNA signal intensity to nuclearly encoded ATP5A signal intensity is also significantly decreased in *actβ* mutants (Fig 5C,D,F). Since mtDNA is found in the mitochondrial matrix and ATP5A is an inner mitochondrial membrane protein, the DNA is adjacent to, but not overlapping with ATP5A (Fig 5C,D, overlay).

**Figure 5:**
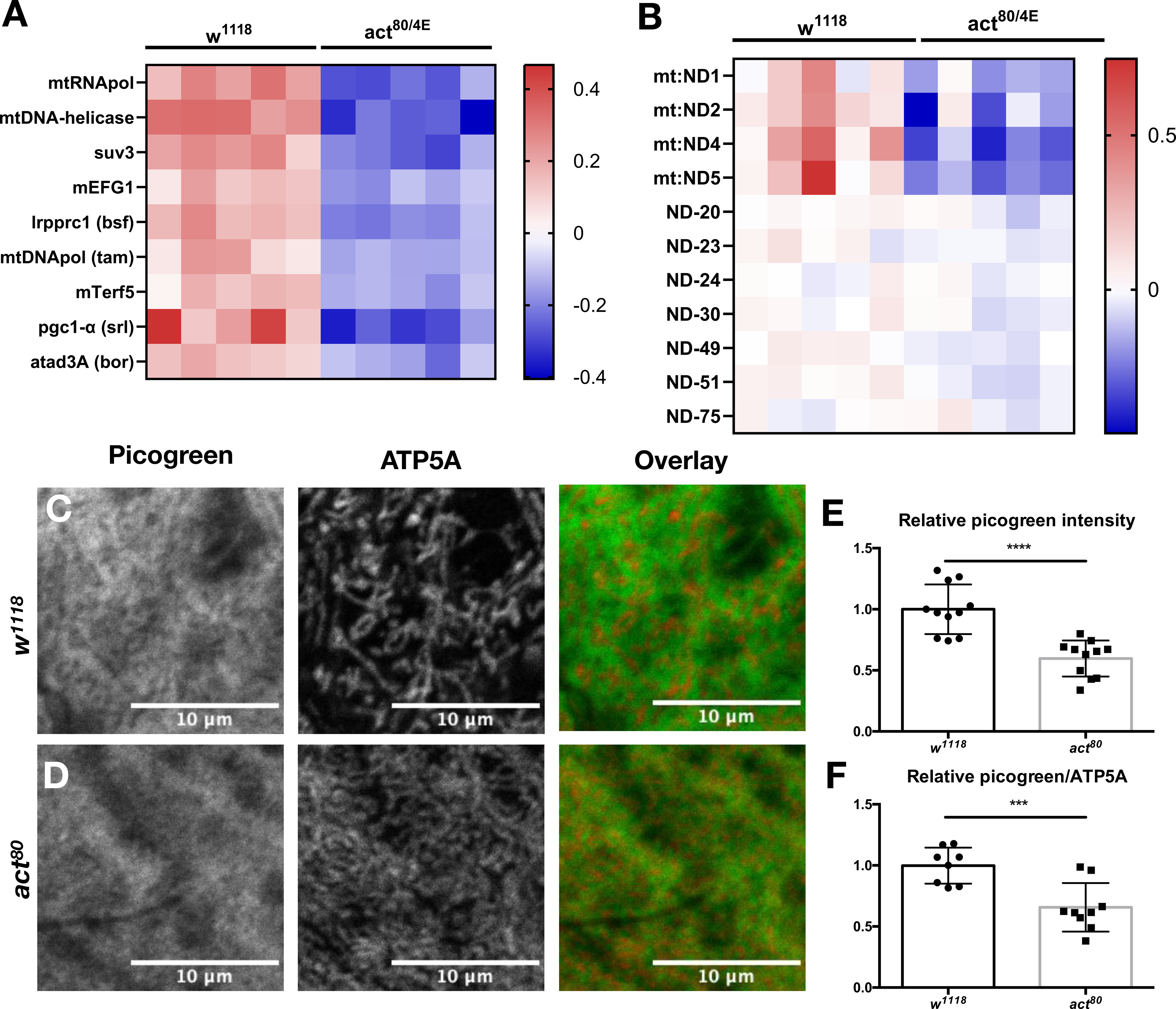
Actβ positively regulates expression of nuclearly encoded factors required for mtDNA maintenance and expression of mitochondrially encoded genes. RNA-seq analysis of control (*w^1118^*) and transheterozygous null *actβ* mutants (*act^80/4E^)* early L3 larvae. Data is plotted as row averages of reads per million, centered and scaled as indicated in the panel. Each column represents pooled RNA from 8 larvae. Biological replicates are shown in the same order in both heat maps. A) p<.0001 for all genes. B) p<.03 for all mitochondrially encoded subunits. Subunits with fewer than 25 reads per million were excluded. Co-staining of early L3 larvae skeletal muscle with ATP5A (nuclearly encoded mitochondrial protein) and picogreen (depicting mtDNA) of C) control (*unc13/+*) and D) homozygous null *actβ* mutants (*act^80^*). E) Quantification of picogreen intensity F) Quantification of the ratio of intensity of picogreen to ATP4A. Each data point represents one animal. ***=p<0.001, ****=p<0.0001

### Knockdown of mtRNA polymerase/PolrMT in muscle results in decreased ratios of mtDNA to nuclear encoded mitochondrial protein and condensed mitochondrial structure

Having identified Actβ as a positive regulator of mtRNA polymerase expression (Fig 5A), we tested whether knocking down *mtRNApol* in the muscle could recapitulate phenotypes observed in *actβ* mutants. We first examined the ratio of total genomic mtDNA to nDNA in the muscle enriched carcass (this includes the epidermis, oenocytes and motor neuron axons which are not expressing RNAi directed at *mtRNApol*) and found a reduction in the ratio of total genomic mtDNA to nDNA (Fig 6A). In agreement with this result, knockdown of *mtRNApol* results in a decreased picogreen (mtDNA) intensity (Fig 6B,C, first panel, D). Additionally, the ratio of picogreen intensity to intensity of nuclearly encoded mitochondrial protein ATP5A is reduced (Fig 6B,C,E) and mitochondrial morphology exhibits a more condensed mitochondrial structure, with shorter rod length similar to that observed in *actβ* mutants (Fig 3E-G). We conclude that the mitochondrial abnormalities observed upon loss of Actβ, including decreased mtDNA, decreased ratio of mtDNA to mitochondrial protein and decreased mitochondrial rod length are likely a result of decreased mtDNA expression resulting from decreased expression of mtRNApol/PolrMT as well as other nulcearly encoded factors involved in mtDNA expression.

**Figure 6:**
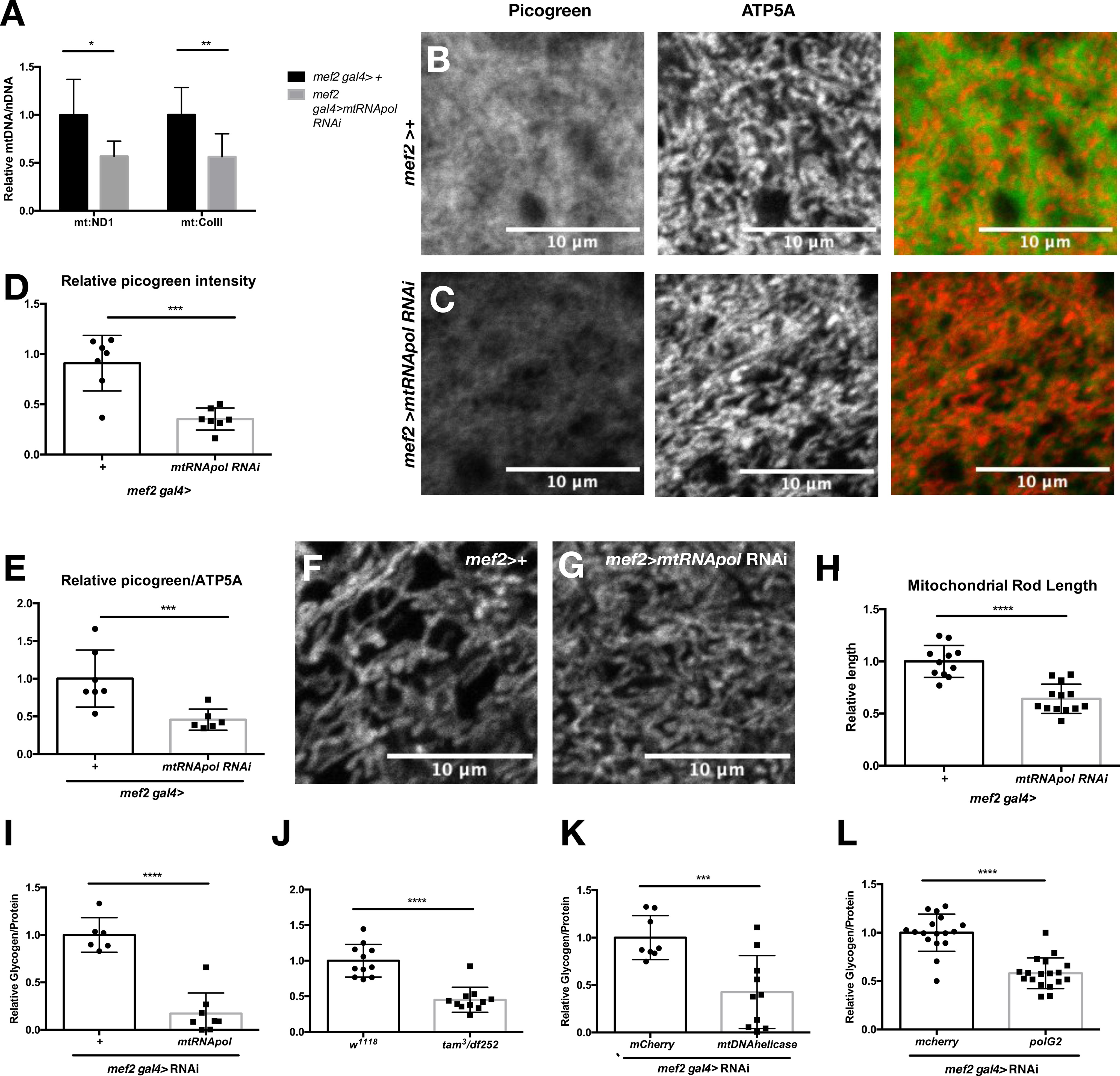
Muscle specific knockdown of *mtRNApol/polrMT* results in decreased mtDNA, mitochondrial rod length and glycogen levels. A) q-PCR Quantification of the ratio of mtDNA (*mt:ND1* and *mt:CoI*) to nDNA (*rpl23*) in the muscle enriched carcass of control (*mef2 gal4 >+*) and muscle specific knockdown of *mtRNApol* (*mef2 gal4>mtRNApol* RNAi) early L3 larvae. N=6 biological replicates consisting of 5 animals each for all conditions. Co-staining of early L3 larvae skeletal muscle with ATP5A (nuclearly encoded mitochondrial protein) and picogreen (depicting mtDNA) in B) control (*mef2 gal4 >+*) and C) muscle specific knockdown of *mtRNApol* (*mef2 gal4>mtRNApol* RNAi). Quantification of D)picogreen intensity and E) the ratio of intensity of picogreen to ATP5A. Each data point represents one animal. Early L3 larvae skeletal muscle stained with ATP5A to show mitochondrial morphology of F) control (*mef2 gal4 >+*) and G) muscle specific knockdown of *mtRNApol* (*mef2 gal4>mtRNApol* RNAi) H) Quantification of mitochondrial rod length. Quantification of total glycogen/protein in early L3 larvae of I) control (*mef2 gal4 >+*) and muscle specific knockdown of *mtRNApol* (*mef2 gal4>mtRNApol* RNAi) J) control (*w^1118^*), and transheterozygous hypomorphic mutants of *tam/mtDNApol* (*tam^3/df252^*), K) control (*mef2 gal4 >+*) and muscle specific knockdown of *mtDNA helicase/ twinkle* (*mef2 gal4> mtDNA helicase* RNAi) and L) control (*mef2 gal4 >+*) and muscle specific knockdown of *polG2* (accessory subunit of mtDNApol) (*mef2 gal4> polG2* RNAi). * = p<0.05, **=p<0.01, ***=p<0.001, ****=p<0.0001

### Decreased levels of nuclearly encoded mtDNA expression factors results in decreased glycogen levels during development

Having shown that Actβ positively regulates both glycogen and nuclearly encoded factors required for mtDNA expression, we next investigated whether decreased expression of proteins involved in mtDNA expression, results in decreased glycogen levels. Knocking down *mtRNApol* in the muscle results in greater than 80% reduction in whole animal glycogen levels (Fig 6H) as well as a slight decrease in glucose levels (Fig S5A). Similarly, a hypomorphic allele of mtDNA polymerase/PolG1 (*tam*), in combination with a deficiency including the *tam* locus, leads to a 60% reduction in glycogen levels (Fig 6I) and a reduction in glucose levels (Fig S5B). Finally, knockdown of *mtDNA helicase/Twinkle* (Fig 6J) or *polG2* (Fig 6K), an accessory subunit of mtDNA polymerase, in muscle produces a 40-60% reduction in glycogen levels, with no change in glucose levels (Fig S5C,D). We conclude that several key nuclearly encoded factors involved in mtDNA expression act as positive regulators of glycogen levels during *Drosophila* development.

### Expressing a constitutively active form of Baboon in muscle of *actβ* mutants, increases mtDNA levels, mitochondrial rod length and glycogen levels

Having seen that knockdown of *actβ* in motor neurons (Fig 1J) or knockdown of *baboB* in muscle (Fig 1K) results in decreased glycogen levels, we reasoned that if Actβ signals directly to the muscle via activation of Baboon, then we should be able to rescue phenotypes resulting from loss of Actβ by expressing a constitutively active form of Baboon (*baboCA*) in muscle of *actβ* mutants. We found that this is indeed the case as expression of constitutively active Baboon in the muscle of an *actβ* mutant increases the ratio of total genomic mtDNA to nDNA in the muscle enriched carcass by 50% (Fig 7A) and increases the ratio of mtDNA (as visualized with picogreen) to mitochondrial protein content (ATP5A) (Fig 7B-D). We also observed an increase in rod length (Fig 7E-G). These data suggest that Actβ signals through Baboon in muscle to regulate mtDNA levels and corresponding mitochondrial morphology.

**Figure 7:**
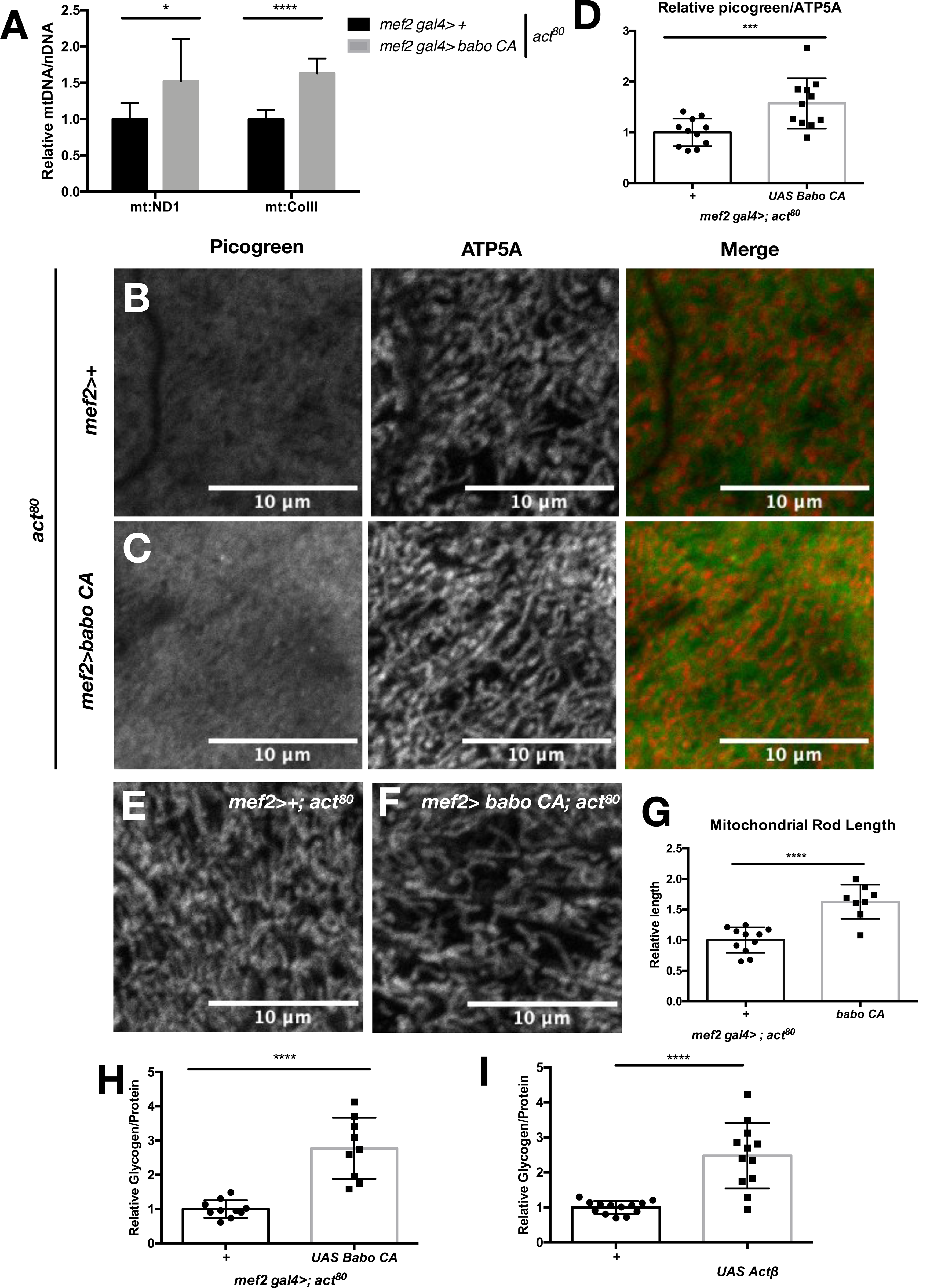
Restoring TGFβ/Activin signaling in muscle of *actβ* mutants results in increased mtDNA, mitochondrial rod length and glycogen levels. A) q-PCR Quantification of the ratio of mtDNA (*mt:ND1* and *mt:CoI*) to nDNA (*rpl23*) in the muscle enriched carcass of early L3 larvae of *actβ* mutants (*mef2 gal4 >+;act80*) and muscle specific activation of the TGFβ/Activin type I receptor Baboon in the *actβ* mutant background (*mef2 gal4>baboCA; act80*). N=7 biological replicates consisting of 5 animals each for all conditions. Co-staining of early L3 larvae skeletal muscle with ATP5A (nuclearly encoded mitochondrial protein) and picogreen (depicting mtDNA) of *actβ* mutants (*mef2 gal4 >+;act80*) and muscle specific activation of the TGFβ/Activin type I receptor Baboon in the *actβ* mutant background (*mef2 gal4>baboCA; act80*). D) Quantification of the ratio of intensity of picogreen to ATP4A. Each data point represents one animal. Early L3 larvae skeletal muscle stained with ATP5A to show mitochondrial morphology of E) *actβ* mutants (*mef2 gal4 >+;act80*) and F) muscle specific activation of the TGFβ/Activin type I receptor Baboon in the *actβ* mutant background (*mef2 gal4>baboCA; act80*)) g) Quantification of mitochondrial rod length. Quantification of total glycogen/protein in early L3 larvae of H) *actβ* mutants (*mef2 gal4 >+;act80*) and muscle specific activation of the TGFβ/Activin type I receptor Baboon in the *actβ* mutant background (*mef2 gal4>baboCA; act80*), I) *actβ* mutants (*ok372 gal4 >+;act80*) and motor neuron specific expression of Actβ in the *actβ* mutant background (*ok372 gal4 >actβ;act80*). In all panels each data point represents 5 individual animals. All experiments were repeated a minimum of 3 times. * = p<0.05, **=p<0.01, ***=p<0.001, ****=p<0.0001

We next investigated whether this manipulation could also increase glycogen levels in *actβ* mutants. Indeed, such expression results in a 2.5-fold increase in glycogen in *actβ* mutants (Fig 7H). Similarly, resupplying Actβ in motor neurons of *actβ* mutants increases glycogen levels by 2.5-fold (Fig 7I). Therefore, we conclude that neuronally derived Actβ positively regulates mtDNA content, mitochondrial rod length and glycogen levels by directly activating TGFβ/Activin signaling in muscle. We hypothesize that the effects Actβ has on mitochondria and glycogen are via regulating the expression levels of several key nuclearly encoded factors involved in mtDNA expression, as knockdown of *mtRNApol/polrMT* in muscle mimics the phenotypes of *actβ* mutants.

## Discussion

Maintaining metabolic homeostasis requires careful coordination of energy usage versus storage at both the organismal and cellular level. Physiological homeostasis is maintained in part by cell signaling pathways. Here we demonstrate that loss and gain of function in *Drosophila* Actβ signal reception in muscles leads to decreased and increased levels of glycogen stores respectively. At the cellular level we identify mitochondria as central regulators of glycogen homeostasis and the likely target in this regulatory circuit. Loss of Actβ leads to a decrease in nuclearly encoded factors involved mtDNA replication, transcription and translation (collectively mtDNA expression) resulting in a concurrent reduction in total genomic mtDNA content. When these same components are reduced through RNAi knockdown in wildtype muscle, we find a similar reduction in total mtDNA levels and a coincident reduction in glycogen content. We conclude that Actβ and TGFβ/Activin signaling provides an unrecognized role in maintaining metabolic homeostasis by regulating glycogen homeostasis and mtDNA expression. This demonstrates that regulation of these processes goes beyond a single organ system and requires communication between two organ systems.

Several previous reports in both *Drosophila* and mammals have linked TGFβ/Activin signaling to carbohydrate homeostasis^35–37,41^. In *Drosophila* the TGFβ/Activin family member, Dawdle, has been shown to be important for regulating genes involved in carbohydrate digestion^41,62^ nuclearly encoded mitochondrial ETC subunits and to be a negative regulator of glycogen in the fat body ^41^. While this latter observation may initially sound counterintuitive since both ligands activate the dSmad2/Smox signal transduction pathway, we believe the production of opposing phenotypes is the result of tissue specific ligand and receptor isoform expression. Dawdle signals exclusively through Babo C^39^, which is the major isoform in fat body^44^, while Actβ signals primarily through the B isoform that is specifically enriched in muscles^40^. Interestingly, Actβ is also produced in intestinal enteroendocrine cells and in response to a high sugar diet this source of Actβ has been implicated to signal through Babo A to augment fat body AkhR (the glucoagon like receptor) expression leading to an increased glycemic index^35^. However, under normal growth conditions we see no Babo A in fat body^44^, although we did observe lower free glucose levels in mid-L3 *actβ* mutants (Fig 1I). Additionally, we did find not find a role for AkhR in regulating glycogen homeostasis (Fig S2F). The explanation for these discrepancies remains to be determined. A separate report has found a role for Actβ in communicating mitochondrial stress from muscle to fat body^63^. This is a very interesting finding and suggests a feedback loop may exist between Actβ and mitochondria.

In mammals, knockout of Follistatin-like 3 (FSTL-3), an Activin and Myostatin antagonist, in mice leads to decreased glycogen stores as well as increased sugar tolerance^64^. This suggests that mammalian Activin and Myostatin function similarly to *Drosophila* Dawdle and negatively regulate glycogen accumulation. On the contrary, Smad3 knockout mice are protected from high fat diet induced hyperglycemia^65,66^. Similarly, hypomorphic *inhbaB* mice, an Actβ homolog, are protected from lipid and glucose accumulation in response to a high fat diet^67^. While we did not test carbohydrate accumulation on a high fat diet, we did look at a high sugar diet and did not see any carbohydrate accumulation in *actβ* mutants (Fig S2G) indicating that the role of Actβ/Smad3/InhbaB in regulating carbohydrate homeostasis may be conserved from *Drosophila* to mammals. As in *Drosophila,* the opposing effects of TGFβ/Activin signaling in the above studies may result from tissue specific differences in activating TGFβ/Activin signaling. Except for the study involving InhabB hypomorphic mice, where the effect on metabolic homeostasis was mapped to regulation of uncoupling proteins^67^, the precise mechanism by which TGFβ/Activin signaling affects cellular metabolism was not investigated.

To explore the role of glucose catabolism in regulating glycogen levels during *Drosophila* development, we undertook a candidate screen in otherwise wildtype flies and measured whole organism glycogen levels (Fig 3). We were initially surprised to find that blocking individual steps in glycolysis and gluconeogenesis had no effect on glycogen homeostasis, however, we speculate that this may be due to the redundancy of some glycolytic enzymes with those of the pentose phosphate pathways and/or the ability of glucose derivatives to enter mitochondria via the glycerol 3 phosphate shunt. The fact that we did not see any changes in glycogen levels when blocking individual steps in glycolysis, a key process in glucose catabolism, demonstrates the robustness of glycogen homeostasis, as might be expected from its importance in health^12,14,15,68^ and development^19,46^. A previous study in *Drosophila* found a role for individual glycolytic genes regulating circulating glucose levels, however this study used older animals^55^, and did not investigate whole animal glycogen content.

We found that limiting glucose derivatives from entering mitochondria resulted in a modest effect on glycogen levels (Fig 3A, G-J). However, this affect was much smaller than the effect on glycogen storage observed with manipulations in TGFβ/Activin signaling (Fig 1). This suggests that the process by which TGFβ/Activin signaling affects glycogen homeostasis is likely independent of basic glucose catabolism. Moreover, it underscores the importance of this pathway in regulating the otherwise very robust process of maintaining glycogen levels. Indeed, we were unable to restore glycogen levels in *actβ* mutants by blocking cytoplasmic glucose catabolism or entry of glucose derivatives into mitochondria (Fig S3). The striking result from our screen was that disrupting oxidative phosphorylation (via knock down of CI or CIV subunits) resulted in a dramatic reduction in glycogen levels (Fig 3K), demonstrating the importance of mitochondrial ETC in maintaining glycogen homeostasis during *Drosophila* development. The role of the ETC in maintaining glycogen levels is not limited to *Drosophila,* cell lines expressing a human mutation in ND75 (CI) show decreased glycogen levels^69^. We speculate the disrupting oxidative phosphorylation may result in altered glucose flux.

One of the many facets that contributes to efficient oxidative phosphorylation is maintaining the optimal balance of total genomic mtDNA to nDNA^1,7,11^. Despite the importance of maintaining this ratio, little is known about how it is regulated. A genome wide screen in *Drosophila* S2 cells for genes that regulate mtDNA abundance identified surprisingly few genes that result in severe mtDNA loss^70^. Outside of factors involved in mtDNA expression, the main class of genes identified that regulates mtDNA levels was components of CV/ATPsynthase. This regulation was found to occur via an increase in reactive oxygen species that subsequently results in mtDNA loss. In addition, the screen found that knockdown of genes involved in cytosolic translation and components of the proteasome resulted in moderate mtDNA depletion^70^. Whether Actβ signaling also acts to modulate the proteasome remains to be examined. However, it is intriguing to note that loss of Baboon/Smox signaling in muscle does decrease translational capacity of muscles^71^.

One of the key aspects of maintaining an optimal balance of mitochondrial and nuclear genomes are components of nuclearly encoded mtDNA expression machinery, yet how these factors are regulated remains largely unknown. A study in mice found that maintaining optimal expression of mtDNApol/PolG1 in CNS and muscle was dependent on a long-ncRNA found in the enhancer locus. However, attempts to identify specific enhancer elements for mtRNApol/PolrMT and mtDNA helicase/ Twinkle was unsuccessful^72^, leaving the regulation of these crucial factors under basal conditions unexplained. Recently, mtRNApol/PolrMT levels have been shown to be regulated by oncogenic myc, and knockdown of PolrMT results in selective tumor cell apoptosis^73,74^. In addition, knockdown or inhibition of PolrMT in acute myeloid leukemia cells results in decreased cellular proliferation and sensitizes resistant cells to apoptosis suggesting PolrMT may be an attractive target for acute myeloid leukemia treatment^75,76^ Our identification of Actβ as an important direct or indirect positive regulator of mtRNApol/PolrMT and mtDNA helicase/Twinkle offers a new mechanism to consider how expression of PolrMT is regulated under basal conditions and how cells in different tissues coordinate expression of nuclear and mitochondrial ETC components to maintain metabolic homeostasis.

The importance of maintaining mitochondrial homeostasis is demonstrated by the consequences of mitochondrial diseases^1,10,11,69,72^. Many of the symptoms of defective mtDNA expression such as muscle fatigue and increased lactic acidosis are indicative of large-scale changes in metabolic homeostasis. In our study we demonstrate that altering mtDNA expression results in large scale changes in glycogen homeostasis during *Drosophila* development. If this is conserved in mammals it may contribute to the muscle fatigue observed in patients and be caused, in part by, increased lactic acidosis. Several previous reports have indicated that mitochondrial and glycogen homeostasis may be intimately connected. For example, an important function of glycogen is to maintain blood sugar levels during fasting and patients with mitochondrial diseases often present with hypoglycemia^8,77,78^. Additionally, in developing *Drosophila* ovarioles, both glycogen levels and mtDNA increase as ovarioles mature^79^.

In addition to showing that decreases in mtDNA expression factors deplete glycogen (either by loss of Actβ or *mtRNApol* knockdown), we find a role for mtDNA expression in regulating tubular/ rod shaped mitochondria morphology (Fig 4E-J, Fig 6E-G). A relationship between mtRNApol levels and mitochondrial tubularity has also been observed in gonads of developing *C. elegans* in which mitochondrial tubularity increases with increased expression of R-POM-1, *C. elegans* mtRNApol^80^. Furthermore, changes in mitochondrial fission and fusion cycles have been linked to changes in mtDNA levels and mtDNA nucleoid structure. Some, but not all, patients with mutations in fusion proteins (such as MFN2 and OPA1) as well as cytoplasmic fusion adaptor proteins show decreased mtDNA^81^, suggesting that this relationship may be bidirectional. Alterations in mitochondrial morphology have also been shown to affect glycogen homeostasis. In colorectal cancer cells impairing mitochondrial fission results in increased glycogen content resulting from increased expression levels of GYS1, which is required to catalyze the rate limiting step of glycogen synthesis. Increased glycogen in these cells increases tolerance to glucose starvation^82^.

In conclusion, we identify Actβ/TGFβ/Activin signaling as a novel upstream signal that changes mitochondrial structure, mitochondrial genome regulation and glycogen homeostasis. Since this signal originates in neurons and affects muscle, it is apparent that the complexity of maintaining glycogen homeostasis and balancing the ratio of total genomic mtDNA to nDNA via expression of nuclearly encoded factors required for mtDNA expression goes beyond the single cell or even individual tissue level and requires coordination across multiple organ systems to maintain whole animal glycogen and mitochondrial homeostasis.

## Acknowledgements

The authors wish to thank Tony Bretscher, Myung-Jun Kim and Tom Neufeld for comments on the manuscript and Matt Sieber and Aidan Peterson for valuable advice throughout the course of this study. This work was funded, in part, by. NIH K12 GM119955 to H.B and R35 GM118029-08 to M.B.O.

## Author contributions

H.B. conducted experiments. H.B and M.B.O conceived and designed the study, interpreted the data, and wrote the manuscript.

---

**Supplemental Figure 1:**

A) Quantification of total glucose/ protein for animals shown in Fig 1A. B) Quantification of 2-NBDG intensity in muscle of control (*w^1118^*) and *actβ* mutants (*act^80^*) early L3 larvae. Representative images if 2-NDBG in four skeletal muscle segments of C) control (*w^1118^*) and D) *actβ* mutants (*act^80^*) early L3 larvae. * = p<0.05, **=p<0.01

**Supplemental Figure 2:**

A, C-E) Quantification of total glycogen/protein in early L3 larvae raised on apple juice plates supplemented with yeast paste (60% w/v in water) of designated genotypes. B) q-RTPCR of *glyS* and *glyP* levels in control (*w^1118^*) and *actβ* mutants (*act^80^*). N=4 biological replicates consisting of 5 early L3 f) Quantification of glycogen/protein in in control (*w^1118^*) and *actβ* mutants (*act^80^*) early L3 larvae raised on a diet consisting of 5 parts yeast and 30 parts dextrose (5Y30Dex) w/v. In all panels with glycogen quantification, all experiments were repeated at least 3 times and each data point consists of 5 animals. ****=p<0.0001

**Supplemental Figure 3:**

A-G) Quantification of total glycogen/protein in early L3 larvae raised on apple juice plates supplemented with yeast paste (60% w/v in water) of designated genotypes. All experiments were repeated at least 2 times and each data point consists of 5 animals., **=p<0.01, ***=p<0.001, ****=p<0.0001

**Supplemental Figure 4:**

A) Western blot showing expression level of nuclearly encoded mitochondrial protein ND30/ NDFSU3 in early L3 larvae of control (*w^1118^*), control (*unc13/+*), *actβ* heterozygotes (*act^80^/+*), and homozygous null *actβ* mutants (*act^80^*). All lanes represent 5 animals and 20μg of protein was loaded. D) Quantification of ND30 expression levels across 3 independent western blots. Image depicting mitochondrial rod length quantification in C) control (*unc13/+*) and D) *actβ* mutants (*act^80^*). Each line drawn represents one of five rods measured in each image. E) Length of each rod in images. Lengths were averaged to create one biological replicate.

**Supplemental Figure 5:**

A-D) Quantification of total glucose/ protein for animals shown in Fig 6 I-L. * = p<0.05, **=p<0.01.

## Basic fly husbandry

Fly lines were maintained on standard cornmeal-yeast-agar medium at 25°C. Unless otherwise noted for all experiments eggs were collected for a three-hour period on an apple juice/agar plate supplemented with yeast paste (approximately 2 parts yeast to 1 part water). Post hatching larvae were transferred to fresh apple juice/ yeast plates at low density and raised until early L3 (approximately 68 hours after egg lay). For high sugar diet, food consisted of 5% yeast and 30% dextrose and 0.8% agar w/v in water.

## Glucose and Glycogen Quantification

Five early L3 larvae were homogenized in 100 μl PBS on ice with a hand held tissue homogenizer and centrifuged at 4°C 2.7k RPM for 3 minutes. An aliquot was removed for protein quantification. Samples were then heat shocked in a 70° water bath for 5 minutes then cooled on ice for 5 minutes. Finally, samples were centrifuged at 4°C for 10 minutes at 13k RPM. Resulting lysates were incubated with Glucose Reagent (Sigma) both in the presence and absence of amyloglucosidase (Sigma) (to digest glycogen) at 37°C for 30 minutes. Absorbance was read at 540nm on a Tecan10M and resulting absorbances were compared to a standard. Glycogen content was quantified by subtracting the absorbance with and without amyloglucosidase.

## Protein Quantification

Five early L3 larvae were homogenized in 100 μl PBS on ice with a hand help tissue homogenizer and centrifuged at 4°C 2.7k RPM for 3 minutes. Lysates were incubated with Pierce BCA Protein Reagent and resulting absorbances were read at 562 nm on a Tecan10M and compared to a protein standard.

## TAG Quantification

Five early L3 larvae were homogenized in 500 μl 5% Nonident P40 on ice with a handheld tissue homogenizer. Lipids were quantified per supplier protocol. Briefly, samples were heated to 80°C 2x and cooled on ice and then spun at 14k rpm for 2 minutes. 15 μl of resulting lysate was incubated in assay bu\er containing a lipase (provided) at room temperature for 20 minutes in the dark. 25μl of assay bu\er Triglyceride probe and enzyme (provided) were added and plate was incubated for 1 minute at room temp. The plate was read a 570nm on a Tecan10M. Lipid concentrations were normalized to protein.

## Periodic Acid Schi<s Reagent Staining

Tissues were dissected in HL3 bu\er and fixed for 30 minutes at RT in 4% formadelhyde. Tissues were than washed with ddH2O and incubated in 0.15% Periodic Acid (1:4 dilution of stock with water) at RT for 15’ on a rotator. Samples were washed with ddH2O and incubated in Schi\’s reagent at RT for 2 minutes on a rotator. Reagent was removed and replaced with stop solution (0.25M HCl, 3% Borax/sodium borate) for 3 minutes at RT on a rotator. Samples were then washed with ddH2O, co-stained with DAPI and imaged immediately.

## 2-NBDG uptake Assay

Larval muscle filets were dissected in PBS and incubated in 2mM 2-NBDG (2-[*N*-(7-nitrobenz-2-oxa-1,3-diazol-4-yl)amino]-2-deoxyglucose) and Hoecsht for 45 minutes at RT on a rotator. Filets were washed two times with PBS and mounted (without fixation) on a piece of double-sided sticky tape and imaged immediately. Due to the variability between muscle segments, signal intensity was measure for four adjacent segments in muscles 6 and 7 and averaged for each biological replicate.

## Starvation Assay

96 well plates were prepared with 200μl of PBS + 0.8% agar. Individual early L3 larvae were washed in PBS and transferred individually to di\erent wells (to prevent cannibalization). Plates were sealed with packing tape and a fine needle was used to poke three holes/ well in the tape. Plates were incubated at 25°C for 72 hours and the surviving larvae were quantified.

## RNA extraction for transcriptomics and qRT PCR

Eight early L3 larvae were homogenized in 500 μl Trizol. 100 μl of cholorform was added and samples were shaken and then incubated at room temperature for 2 minutes. Samples were then spun at 11k for 15’ at RT and the aqueous phase was removed. Equal volumes of absolute ethanol was added and applied to a Zymo RNAmicroprep column. RNA was purified per manufactures protocol. RNA concentrations were quantified using a nanodrop.

## cDNA synthesis and qRT PCR

cDNA was synthesized from 2 μg of RNA using SuperScript III First Strand synthesis kit as per manufactures’ protocol. Resulting cDNA was diluted 1:5 with DNA/RNAase free water and used as a template for qRT PCR. SYBR green I master mix kit was used on a Light Cycler 480 (Roche) and expression levels were normalized to a house keeping gene.

## Mitochondrial DNA quantification

Five early L3 larvae were squished in 50 μl squish bu\er (10mM Tris-Hcl pH 8.0, 1mM EDTA, 25mM NaCl and 0.2mg/ml proteinase K) and incubated at 37°C for 30 minutes, followed by 5 minutes at 95°C to inactivate proteinase K. cDNA was diluted with DNA/RNAase free water and used as a template for q PCR. SYBR green I master mix kit was used on a Light Cycler 480 (Roche) and ratio of mtDNA to nDNA was calculated.

## Immunohistochemistry

Animals were dissected in Ca^2+^-free HL3 bu\er and fixed with 4% formaldehyde at RT for 30 minutes. Muscle filets were then washed with PBS and blocked in PBS+ 0.1% Triton +1% NGS for 30 minutes at RT on a rotator. Tissues were stained with ATP5A (1:250) diluted in blocking solution overnight at 4 degrees on a rotator. Tissues were washed once with PBS and then incubated in blocking solution at RT for 20 minutes on a rotator. Tissues were incubated in secondary antibody (1:1000) and picogreen (1:200) in blocking solution and incubated at RT for 2 hours on a rotator protected from light. Samples were washed once with PBS and mounted in vectashield and imaged immediately.

Images were acquired on a Zeiss LSM 710 confocal microscope equipped with a 20x Plan-Apochromat 20x/0.8 M27 and a 100x alpha Plan-Apochromat 100x/1.46 Oil DIC M27objective. ImageJ was used to quantify the intensity of the picogreen and ATP5A staining in a 200x200 pixel area located in the center of muscle 6 or 7. ImageJ was used to rod length in a 200x200 pixel area located in the center of muscle 6 or 7. For each area the 5 longest distinct rod structures were measured and averaged for each biological replicate.

## Western Blot Analysis

Five early L3 larvae were homogenized in 21 μl of RIPA bu\er supplemented with a cocktail of protease inhibitors and incubated at 4°C for 40 min with on a rotator. Protein concentration of lysates was analyzed and 20 μg of protein was loaded on 4-12% Bis-Tris gel and transferred onto PVDF membrane. The membranes were then blocked with Casein-containing bu\er and incubated with primary antibody at 4°C overnight.

## Transcriptomics

RNAi was extracted from 8 animals/ biological replicate for each genotype. Total RNA (3 μg per sample) was submitted to the University of Minnesota Genomics Center (UMGC) for quality assessment and Illumina next-generation sequencing. 2 x 150bp FastQ paired-end reads for samples (n=30.3 Million average reads per sample) were trimmed using Trimmomatic (v 0.33) enabled with the “-q” option; 3bp sliding-window trimming from 3’ end requiring minimum Q30. Quality control on raw sequence data for each sample was performed with FastQC. Read mapping was performed via Hisat2 (v2.1.0) using the *Drosophila* genome (*Drosophila melanogaster* (BDGP6.28) as a reference. Gene quantification was done via Feature Counts for raw read counts.

