## Supplementary figures and images for "Glycogen homeostasis and mtDNA expression require motor neuron to muscle TGFβ/Activin Signaling in *Drosophila*"

### Supplemental Figure 1

#### Supplemental Figure 1

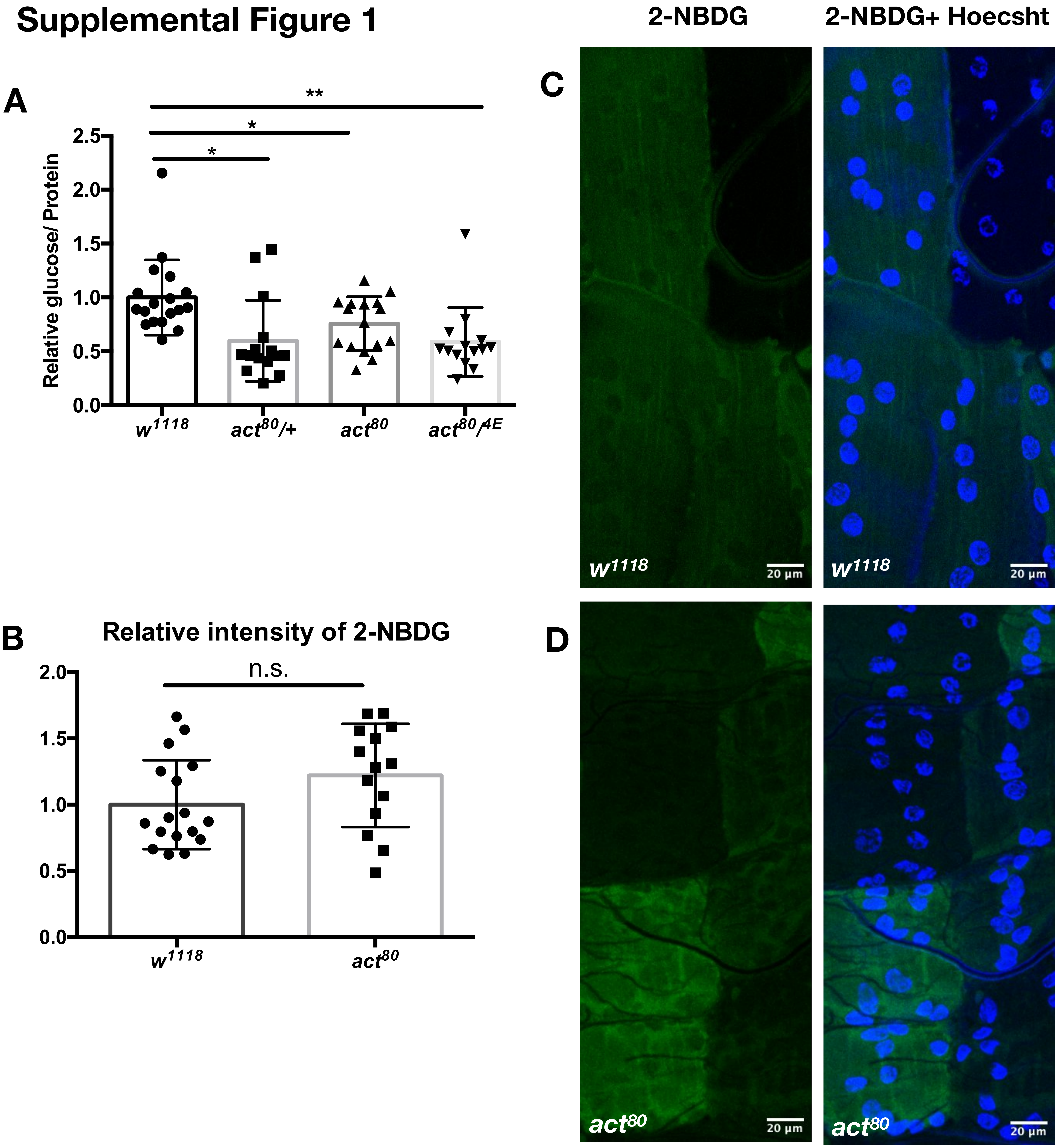

### Supplemental Figure 2

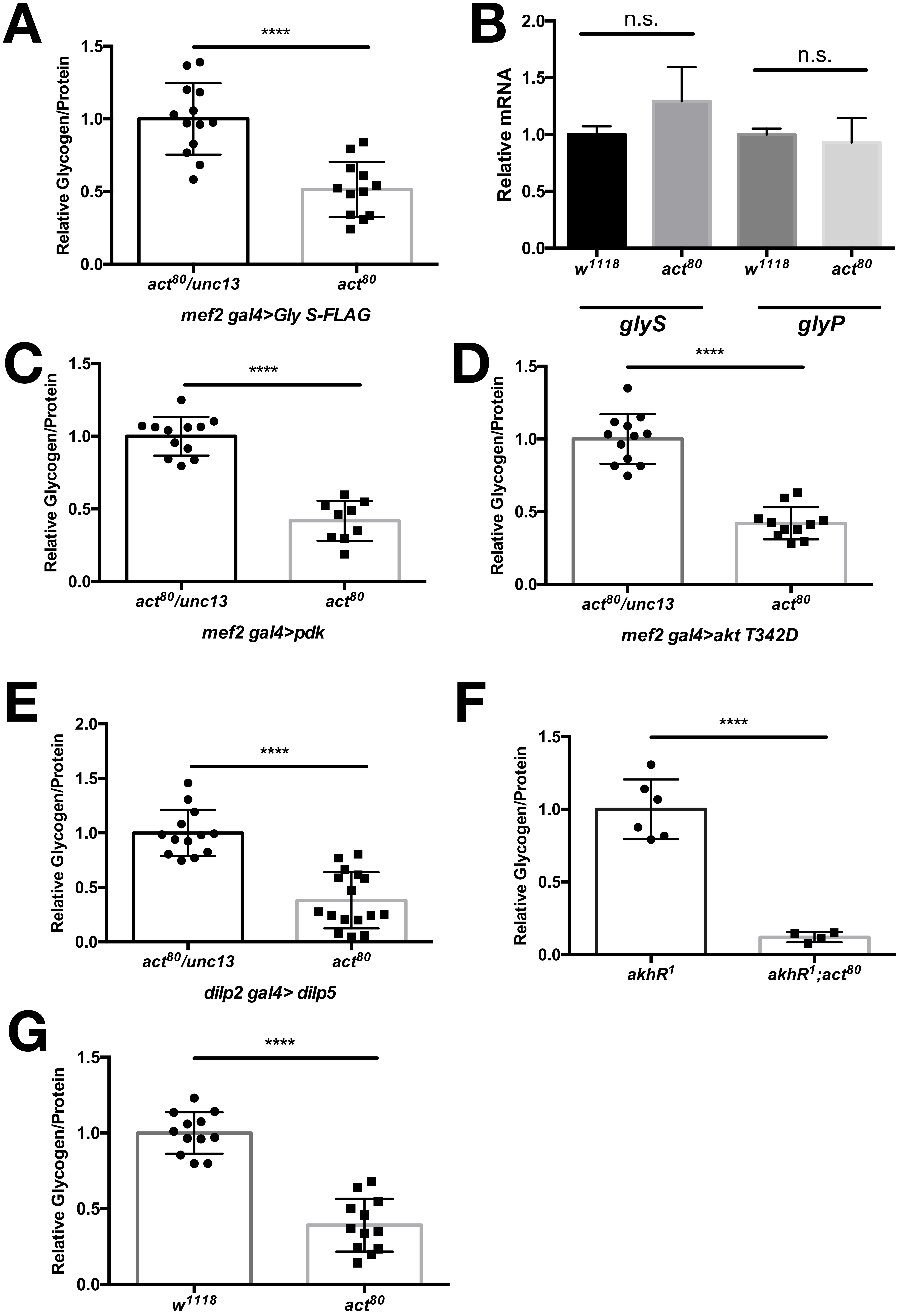

### Supplemental Figure 3

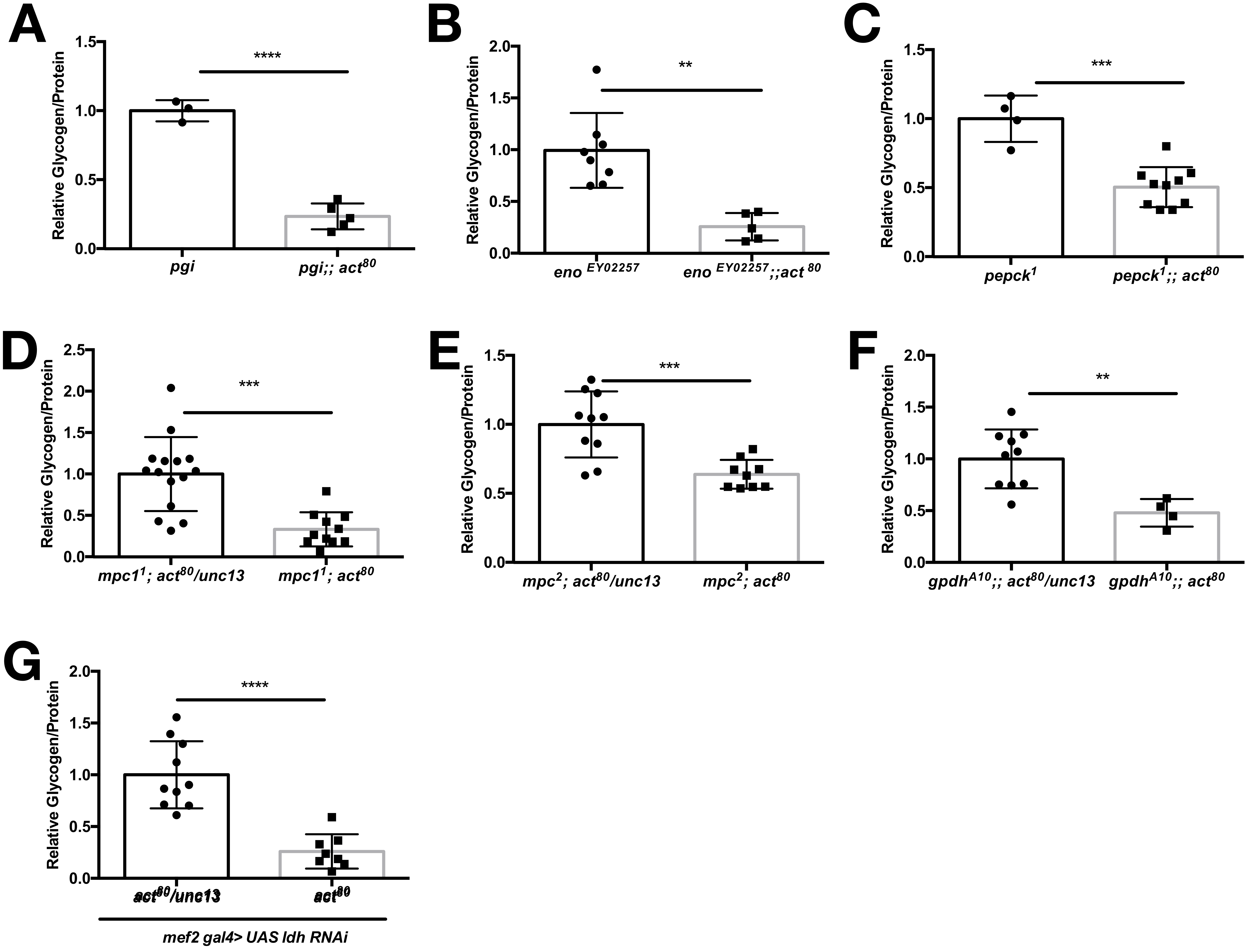

### Supplemental Figure 4

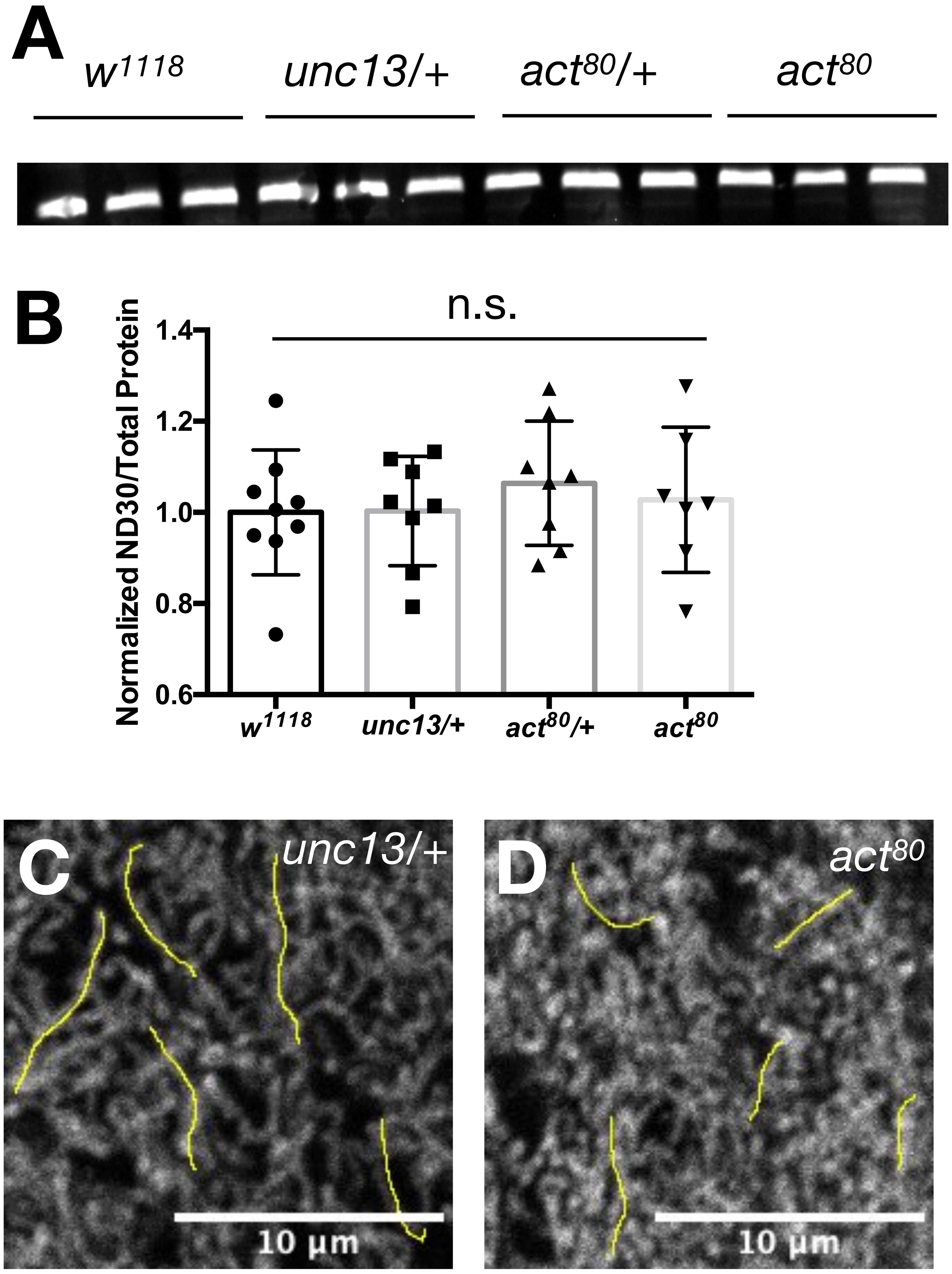


E

| | *unc13/+* | *act*$^{80}$ |
|---|---|---|
| | 7.79 | 3.245 |
| | 7.368 | 5.282 |
| | 5.917 | 3.613 |
| | 5.35 | 3.016 |
| | 6.196 | 4.433 |
| Average | 6.5242 | 3.9178 |

### Supplemental Figure 5

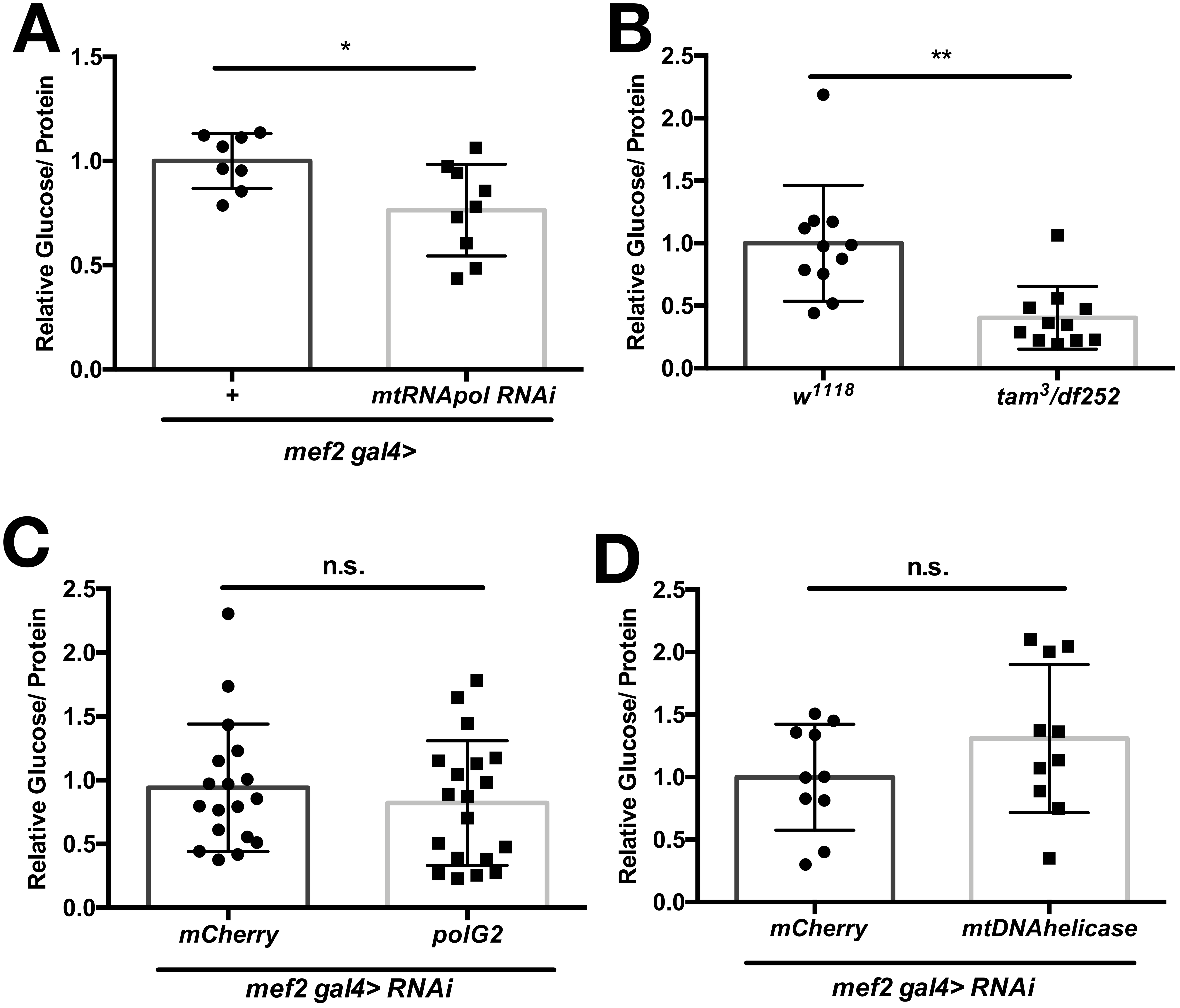
